# Fertilisation of agricultural soils with municipal biosoilds: Part 2- Site properties, environmental factors, and crop identify influence soil bacterial communities more than municipal biosolid application

**DOI:** 10.1101/2023.12.14.571735

**Authors:** Andrew J.C. Blakney, Simon Morvan, Marc Lucotte, Matthieu Moingt, Ariane Charbonneau, Marie Bipfubusa, Emmanuel Gonzalez, Frédéric E. Pitre

## Abstract

Reducing the environmental impact of Canadian field crop agriculture, including the reliance on conventional synthesised fertilisers, are key societal targets for establishing long-term sustainable practices. Municipal biosolids (MSB) are an abundant, residual organic material, rich in phosphate, nitrogen and other oligo-nutrients, that could be used in conjunction with conventional fertilisers to decrease their use. Though MBS have previously been shown to be an effective fertiliser substitute for different crops, including corn and soybean, there remain key knowledge gaps concerning the impact of MBS on the resident soil bacterial communities in agro-ecosystems. We hypothesised that the MBS fertiliser amendment would not significantly impact the structure or function of the soil bacterial communities, nor contribute to the spread of human pathogenic bacteria, in corn or soybean agricultural systems. In field experiments, fertiliser regimes for both crops were amended with MBS, and compared to corn and soybean plots with standard fertiliser treatments. We repeated this across four different agricultural sites in Quebec, over 2021 and 2022. We sampled MBS-treated, and untreated soils, and identified the composition of the soil bacterial communities via 16S rRNA metabarcoding. We found no indication that the MBS fertiliser amendment altered the structure of the soil bacterial communities, but rather that the soil type and crop identities were the most significant factors in structuring the bacterial communities. Moreover, there was no evidence that the MBS-treated soils experienced a shift in functions, nor contributed potential human bacterial pathogens over the two years of our study. Our analysis indicates that not only can MBS function as substitutes for conventional, synthesised fertilisers, but that they also do not disrupt the structure, or function, of the resident soil bacterial communities in the short term. Finally, we suggest that the use of MBS in agro-ecosystems poses no greater concern to the public than existing soil bacterial communities.

**Highlights:**

- Municipal biosolids may represent a sustainable fertiliser substitute
- But, the impact of biosolids on soil bacteria in agricultural fields is unknown
- Using 16S rRNA metabarcoding we analysed community structure and functions
- We found no disruption of soil bacterial communities fertilised with biosolids
- Biosolids are safe, sustainable fertilisers with little impact on soil bacteria

**Graphical Abstract:** 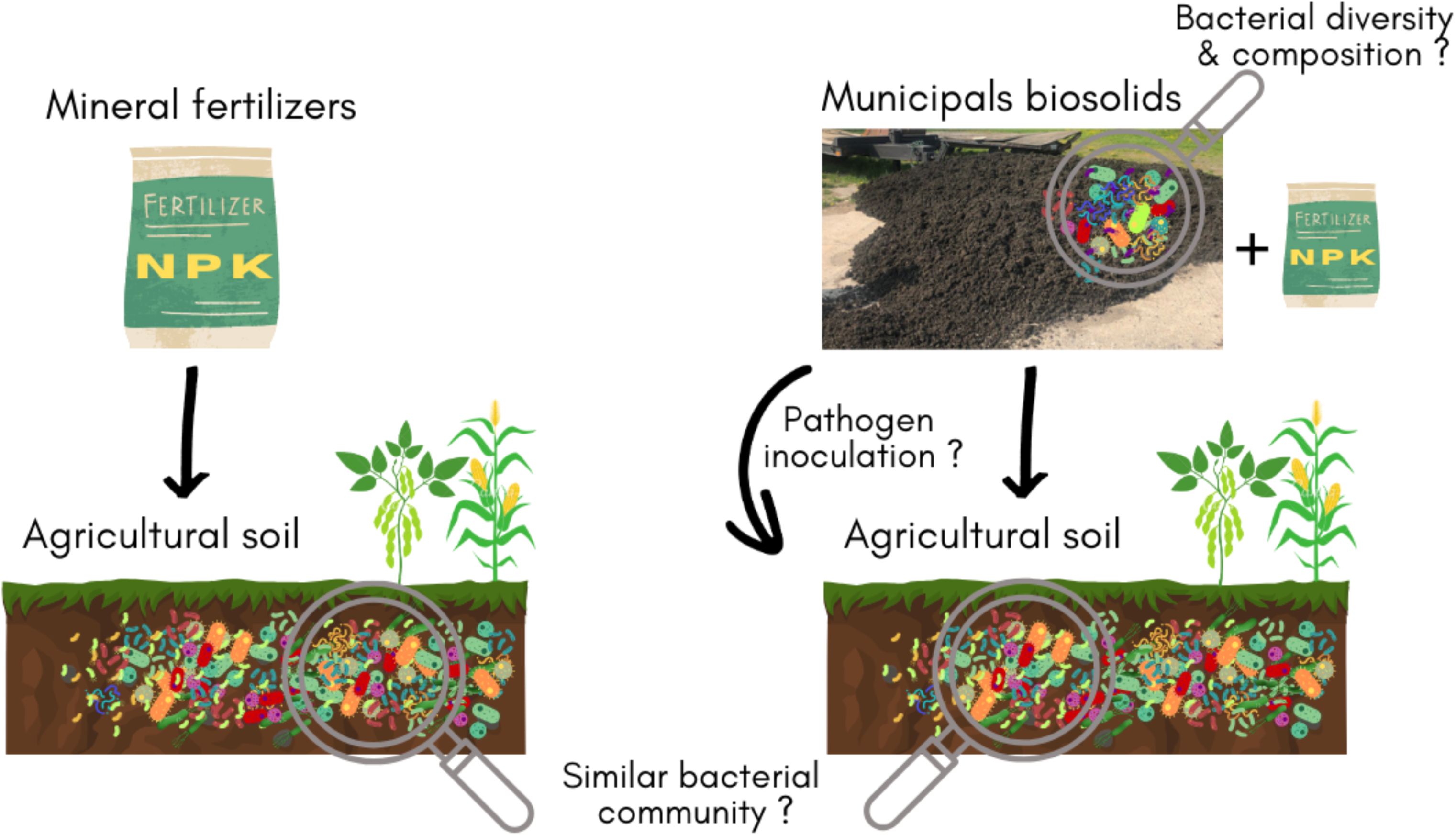

## 1.0 Introduction

Global demand for increased agricultural production is straining our environment and natural resources (Wittwer *et al*., 2021; Halpern *et al*., 2022). The agriculture sector alone accounts for ∼28% of global anthropogenic greenhouse gases, including CO2, CH4, and N2O (Tubiello *et al*., 2015; Crippa *et al*., 2021), and uses 70% of the available fresh water (Rosegrant *et al*., 2009). Furthermore, the majority of anthropogenic nitrogen and phosphorus produced and applied as mineral fertiliser, continue to linger and cycle through the biosphere for years (Keeler *et al*., 2016; Zou *et al*., 2022). This translates to wasted energy in their production and application, and widespread environmental degradation (Keeler *et al*., 2016; Halpern *et al*., 2022; Zou *et al*., 2022). As such, alternatives to synthetic and mineral fertilisers are necessary to reduce the impact of nitrogen and phosphorus inputs in agriculture, including recognizing their social and environmental cost, finding more targeted and efficient uses, and effective substitutes (Keeler *et al*., 2016; Milner & Boyd, 2017; Fargione *et al*., 2018; Agriculture & Agri-Food Canada, 2022; Zou *et al*., 2022).

Municipal biosolids (MBS)^1^ are an abundant, residual organic material, rich in phosphate, nitrogen and other oligo-nutrients. As such, it has been suggested that MBS could be used in conjunction with conventional fertilisers in order to decrease their use (Schlatter *et al*., 2019). As organic fertilisers, MBS ought to release their nutrients progressively, as opposed to the immediate nutrient availability of chemical fertilisers (Schlatter *et al*., 2019). Moreover, several studies have gone on to illustrate the effectiveness of MBS as a fertiliser substitute especially in field crop agriculture (Vasseur *et al.,* 1999; Warman & Termeer, 2005; Gardner *et al*., 2009). In our companion article, Charbonneau *et al*. (2023) test the efficiency of MBS as a fertiliser amendment for corn and soybean in sites around Québec. As a proof-of-concept, we found yields were equal among crops treated with MBS in conjunction with reduced doses of mineral fertilisers compared to those treated only with conventional fertilisers.

However, adding MBS into agricultural systems raises important questions concerning the microbiological impact on, first, the resident soil bacteria, and second, spreading pathogenic bacteria into the food supply-chain. The microbiology of MBS is well documented and does contain more enteric and potentially opportunistic bacterial pathogens than might be typically found in the soil, such as *Escherichia coli*, *Enterococcus* sp., *Yersina* sp, and *Campylobacter* sp., among others (Ryan *et al*., 2009; Walterson *et al*., 2015; Depoorter *et al*., 2016; Scott *et al*., 2022). Although local regulations require that any MBS to be applied as fertiliser be tested for safe levels of coliforms, data is still missing with regard to if the applied MBS introduces, or increases the risk of bacterial pathogens in agricultural soils.

Furthermore, introducing the novel bacteria from MBS into agricultural systems may alter or disrupt the structure and function of the resident soil bacteria (Strickland *et al*., 2009; Walsh *et al*., 2021). These complex soil bacterial communities form over time through the on-going interactions of the existing soil chemistry, the extant microbial community, and any exogenous chemical inputs, including from plant material and their agricultural treatments (Blakney *et al*., 2022). These communities provide critical functions in agro-ecosystems, including nutrient cycling (Richardson *et al*., 2009; Weidner *et al*., 2015; Yu *et al*., 2021), tempering environmental changes (Lau & Lennon, 2012), or plant host stress (Marasco *et al*., 2012; Hou *et al*., 2021), and protecting plant hosts against pathogens (Sikes *et al*., 2009; Mendes *et al*., 2011). Currently, it remains unclear if inoculating agricultural soils with MBS, and the bacteria they contain, may alter the structure and functions of the existing soil bacterial communities.

Here, we address these knowledge gaps concerning the impact of MBS as a fertiliser amendment in agro-ecosystems on soil bacterial communities. We hypothesised that the MBS fertiliser amendment would not significantly impact the structure nor function of the soil bacterial communities in corn or soybean agricultural systems. In our field experiments, corn and soybean plots were amended with either a combination of MBS and reduced doses of mineral fertilizers, or a regular dose of mineral fertiliser. We repeated this across four different agricultural sites in Quebec, over 2021 and 2022. We sampled MBS-treated, and untreated soils, and identified the composition of the soil bacterial communities via 16S rRNA metabarcoding. To evaluate our hypothesis we looked for changes among bacterial community composition, structure, and functional roles, in MBS treated agricultural soils, compared to control plots.

## 2.0 Methods

### 2.1 Site description & experimental design

As described in Charbonneau *et al*. (2023), a field experiment was established in four agricultural sites in Quebec, Canada. Briefly, sites were selected based on similar soil types (silty-clay), and agricultural practices (conventional production methods, direct seeding, corn-soy rotations, and use of glyphosate-based herbicides). At each site, two fields were prepared as a split-plot experimental design replicated in four complete blocks, where each block was randomly divided into municipal biosolid (MBS) treated and untreated plots. In 2021, one field was seeded with Roundup Ready soy and a second field with Roundup Ready corn. In 2022, following the same experimental design, the same fields were rotated, establishing soy-corn and corn-soy rotations over 2021 and 2022 at each site. In total, the experiment contained 128 plots: 2 treatments * 4 blocks * 2 crops * 4 sites * 2 years (See Charbonneau *et al*. (2023) for more details).

### 2.2 Crop management and sampling

As described in Charbonneau *et al*. (2023), crops at all sites were grown and maintained according to the recommendations of the Quebec Reference Center for Agriculture and Agri-food (CRAAQ), standard agricultural practices. MBS was obtained from a regional wastewater treatment plant, and applied to the treatment plots in early May before seeding. The amount of MBS spread was determined according to local agricultural by-laws, which dictates the maximal dose of phosphate application permitted based on the average content of phosphorus measured in the MBS of the regional station between 2019 and 2021 (Parent & Gagné, 2010). Similar cumulative amounts of N-P-K were applied to both treated and untreated plots: mineral fertilisers were used for untreated plots while treated plots were fertilised with a combination of mineral fertilisers and MBS (See Charbonneau *et al*., 2023 for more details).

For the microbial community analysis, sampling occurred at two time points: campaign 1 (C1), the baseline control, sampled in May before the MBS treatment, seeding, and the initial use of glyphosate-based herbicide, and C2 in July, during the growing season after the application of MBS and the final application of glyphosate-based herbicide (See Charbonneau *et al*., 2023 for more details). For each plot, 10 soil cores, ∼5 m apart, were sampled using a manual corer at 0-20 cm depth in the row spacing. All 10 cores were then pooled to form composite samples from each plot. Each sample core was geo-referenced for subsequent sampling campaigns over both 2021 and 2022. From each composite soil sample, 15mL was subsampled to conduct microbial community analysis and placed on ice in coolers, before being stored at the Institut de Recherche en Biologie Végétale, Montréal (QC, Canada) at -80°C awaiting for further processing. We accounted for the use of various agricultural treatments in the downstream metabarcoding data by considering each soil sample and their total complement of particular agricultural treatments as a unit.

### 2.3 DNA extraction from soil and biosolid samples

Total DNA was extracted from all the MBS and soil samples using the NucleoSpin Soil DNA extraction kit (Takara Bio, USA), following the manufacturer’s instructions. The SL1 lysis buffer was used for all extractions, with between 300 and 500 mg of soil or MBS sludge. All extracted DNA samples were qualitatively evaluated using a 0.7 % agarose gel, for 35 minutes at 100 V. DNA concentration was also evaluated using a Nanodrop (Thermo Fisher NanoDrop 2000). Extracts were stored at -20°C until they could be sent for amplification and sequencing by Genome Québec Innovation Center (Montréal, QC, Canada).

### 2.4 16S rRNA gene metabarcode generation and sequencing

The prepared plates of DNA samples were submitted to Génome Québec (Montréal, Québec) for 16S rRNA amplicon generation and sequencing (Bell *et al*., 2016; Lay *et al*., 2018; Blakney *et al*., 2022). The bacterial genome region targeted in this experiment was the V5-V6 region of the 16S ribosomal RNA (rRNA) for amplification by PCR using the forward primer: S-D-Bact-0785-a-S-18, GGMTTAGATACCCBDGTA and reverse primer: S-*-Univ-1100-a-A-15, GGGTYKCGCTCGTT (Klindworth *et al*., 2012). The CS1 (ACACTGACGACATGGTTCTACA) and CS2 (TACGGTAGCAGAGACTTGGTCT) tags were used to add barcodes as well as Illumina adapters i5 and i7 that allow sequence binding to the flow-cell. Sequencing was conducted on an Illumina MiSeq using a paired-end 2 x 300 base-pair method (Illumina, San Diego, CA, USA). We estimated this should provide a mean of 60 000 reads per sample, which is in line with previous studies that describe bacterial communities (Bell *et al*., 2016; Lay *et al*., 2018; Blakney *et al*., 2022).

### 2.5 Estimating ASVs from MiSeq 16S rRNA gene metabarcodes

The 16S rRNA gene amplicons generated by Illumina MiSeq were used to estimate the diversity and composition of the bacterial communities present in each soil and MSB sample. The integrity and totality of the 16S MiSeq data downloaded from Génome Québec was confirmed using their MD5 checksum protocol (Roy *et al*., 2018). Subsequently, all data was managed, and analysed in R (4.0.3 R Core Team, 2020), and plotted using ggplot2 (Wickham, 2016).

The bioinformatic Dada2 pipeline (Callahan *et al*., 2016) was used to process and annotate raw sequence reads. Briefly, primers were removed with Cutadapt (Martin, 2011) version 2.10, reads mapping to PhiX Control were removed, forward and reverse reads were trimmed up to position 180 nt, reads containing uncertain base calls were removed, and after trimming, reads containing more than 2 expected errors were discarded. Reads were denoised and subsequently dereplicated with the dada() function using the parameter *pool = TRUE*. Reads were finally assembled into potential amplicon sequence variants (ASV) and chimeras were removed using Dada2’s removeBimeraDenovo() function. ASVs annotation process used SILVA v132 database (Yilmaz *et al*., 2013; Parks *et al*., 2018) and all non-bacterial ASVs were discarded. All annotations should be considered as putative and interpreted with caution as databases contain errors and are subject to change.

After these filtering steps, we obtained 34 142 different ASVs throughout the 2021 dataset with about 25 000 sequences per soil sample (**Table S1**). For the 2022 dataset, we obtained 7886 different ASVs with around 5400 sequences per soil sample (**Table S1**). Rarefaction curves were generated using the rarecurve() function from the vegan R package (Oksanen *et al*., 2020). The soil samples curves show inflections, indicating that we managed to capture most of the bacterial community. (**Fig. S1**). The MBS samples curves systematically reach a plateau, indicating a more thorough capture of the bacterial community in these samples. Finally, we used the phyloseq R package to streamline data handling and produce alpha diversity and beta diversity plots (McMurdie & Holmes, 2013).

### 2.6 α-diversity and β-diversity of the of the soil bacterial communities

Simpson reciprocal (1-D) and Shannon-Weaver (H) α-diversity indices were computed using the plot_richness() function within the phyloseq package (McMurdie & Holmes, 2013). To assess statistical differences between the different groups, we conducted Kruskal-Wallis tests as well as pairwise Wilcoxon rank-sum test when there were more than two groups. For evaluating the similarity of microbial communities, we performed β-diversity analysis employing the Aitchison distance, which involves transforming sequence abundances through a centred log ratio transformation before computing the Euclidean distance between sites, as described by Quinn *et al*. (2018). This procedure takes into account the compositional nature of sequencing data, and is therefore an appropriate dissimilarity metric (see Gloor *et al*., 2017). Principal component analysis (PCA) ordinations were conducted to visualise the resemblance between samples in terms of their microbial communities. To assess the significance of the observed differences, we performed a PERMANOVA using the adonis2() function from the vegan package with 999 permutations. As PERMANOVA is robust against heterogeneity of dispersion for balanced designs (Anderson & Walsh, 2013), we only report dispersion for the unbalanced. The assumption of homogeneity of dispersion was assessed using the betadisper() and permutest() functions, both from the vegan R package (Oksanen *et al*., 2020). We used a pairwise PERMANOVA post-hoc test to identify which groups were significantly different when necessary. Finally, we quantified how soil texture impacted the soil bacterial community structure with a distance-based redundancy analysis. The percentage of clay, silt and sand that composed each site was transformed with a centered-log ratio, appropriate for compositional data. We modelled the explanatory power of soil texture at both time points (May, C1, and July, C2) in corn and soybean fields from both sampling years, using the rda() function in the vegan package (Oksanen *et al*., 2020). Each model was constrained by soil texture, and tested for significance using PERMANOVAs, as described above.

### 2.7 Identifying potential bacterial pathogens

To address concerns from producers and consumers about the presence of human pathogens in MBS, we determined the abundance of 16S rRNA sequences assigned to genera with known potential human bacterial pathogens, including *Escherichia* (*Enterobacteriaceae*), *Salmonella* (*Enterobacteriaceae*), *Listeria* (*Listeriaceae*), which represent > 80% of cases of agricultural-borne illness (Brennan *et al*., 2022). We also looked for more opportunistic, soil-borne pathogens of clinical interest, such as *Stenotrophomonas* (*Xanthomonadaceae*; Brooke, 2021), and *Pseudomonas* (*Pseudomonadaceae*; Rossi *et al*., 2021). We compared the abundance of these taxa to their corresponding bacterial communities across both sampling years, 2021 and 2022, and both crops.

To further assess the presence of potential bacterial pathogens, we also assigned functional categories to the inferred ASVs using the microeco package (Liu *et al*., 2021) and the FAPROTAX database (Louca *et al*., 2016; Sansupa *et al*., 2021). The relevant functional categories included in this analysis were human pathogens, plant pathogens and animal parasites or symbionts. We then compared the number of ASVs assigned with these pathogenic functions from the MBS to the MBS-treated and non-treated soils in both crops, during both years. Differences between the MBS-treated and non-treated soils were tested using the non-parametric Kruskal-Wallis test.

## 3.0 Results

### 3.1 Relative abundance of bacterial communities in MBS and in soils

The initial composition of the bacterial communities in the MBS and soils from the different corn and soybean agricultural sites was determined. In 2021, out of the 32 phyla identified, the MBS samples were dominated by *Actinobacteriota* 29.5%, *Bacteriodota* 19.0%, and *Firmicutes* 13.7%. More precisely, the top three families found were *Propionibacteriaceae* (*Actinobacteriota* – 22.3%), *Cloacimonadaceae* (*Cloacimonadota* – 9.3%) and *Christensenellaceae* (*Firmicutes* – 6.7%). The 2022 MBS communities differed from 2021 with *Bacteriodota* 45.7%, *Firmicutes* 19.3%, and *Proteobacteria* 14.7% as the top three phyla, while *Actinobacteriota* represented only 3.4% of the relative abundance (**Fig. 1A & S2A**). The *Bacteriodota* sequences mainly belonged to the *Dysgonomonadaceae* (21.2%) and *Rikenellaceae* (14.7%) families, while the largest family in the *Firmicutes* was the *Christensenellaceae* (6.1%). Proteobacteria ASVs were mostly classified as *Pseudomonadaceae* (11.3%).

**Figure 1.**
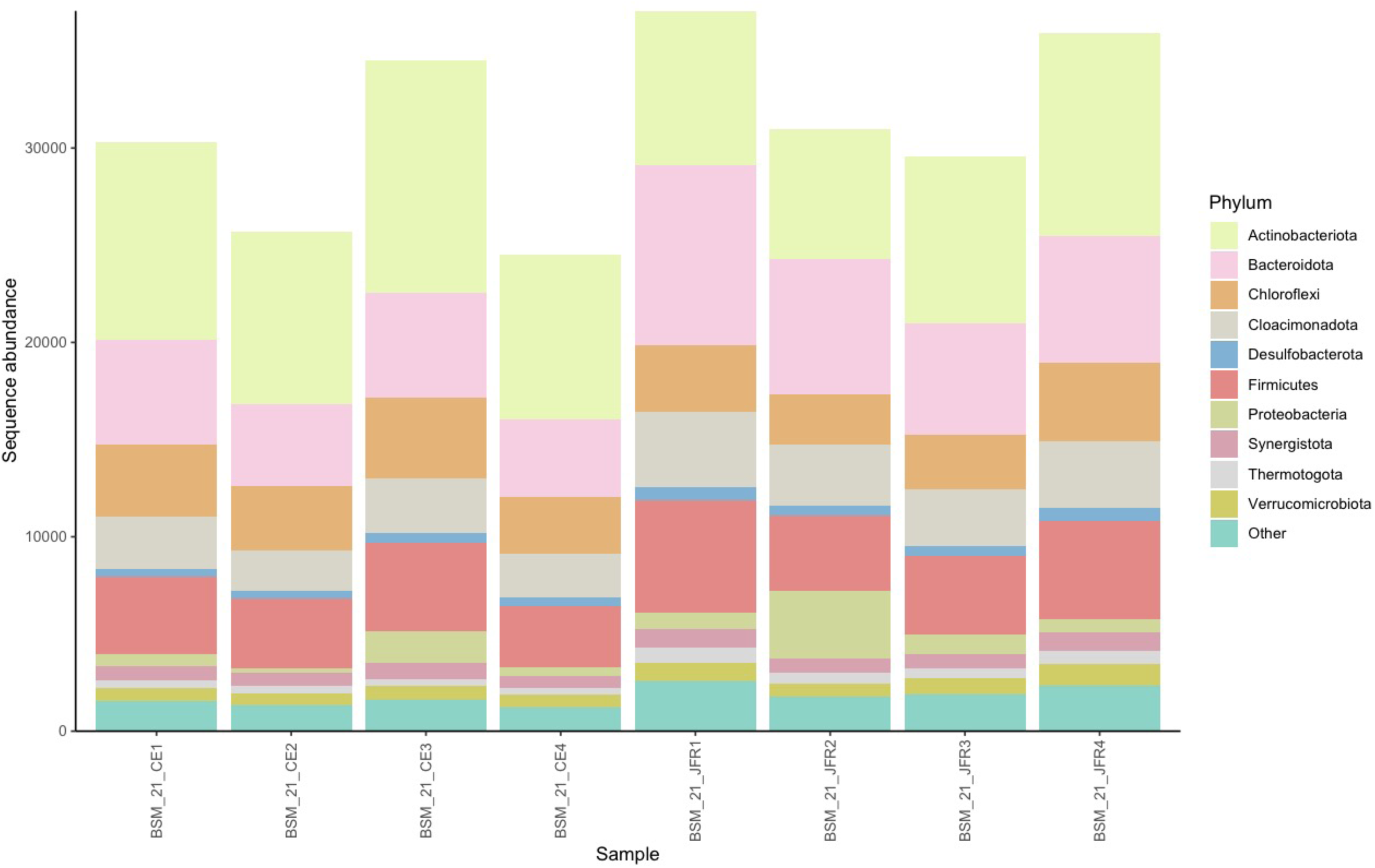
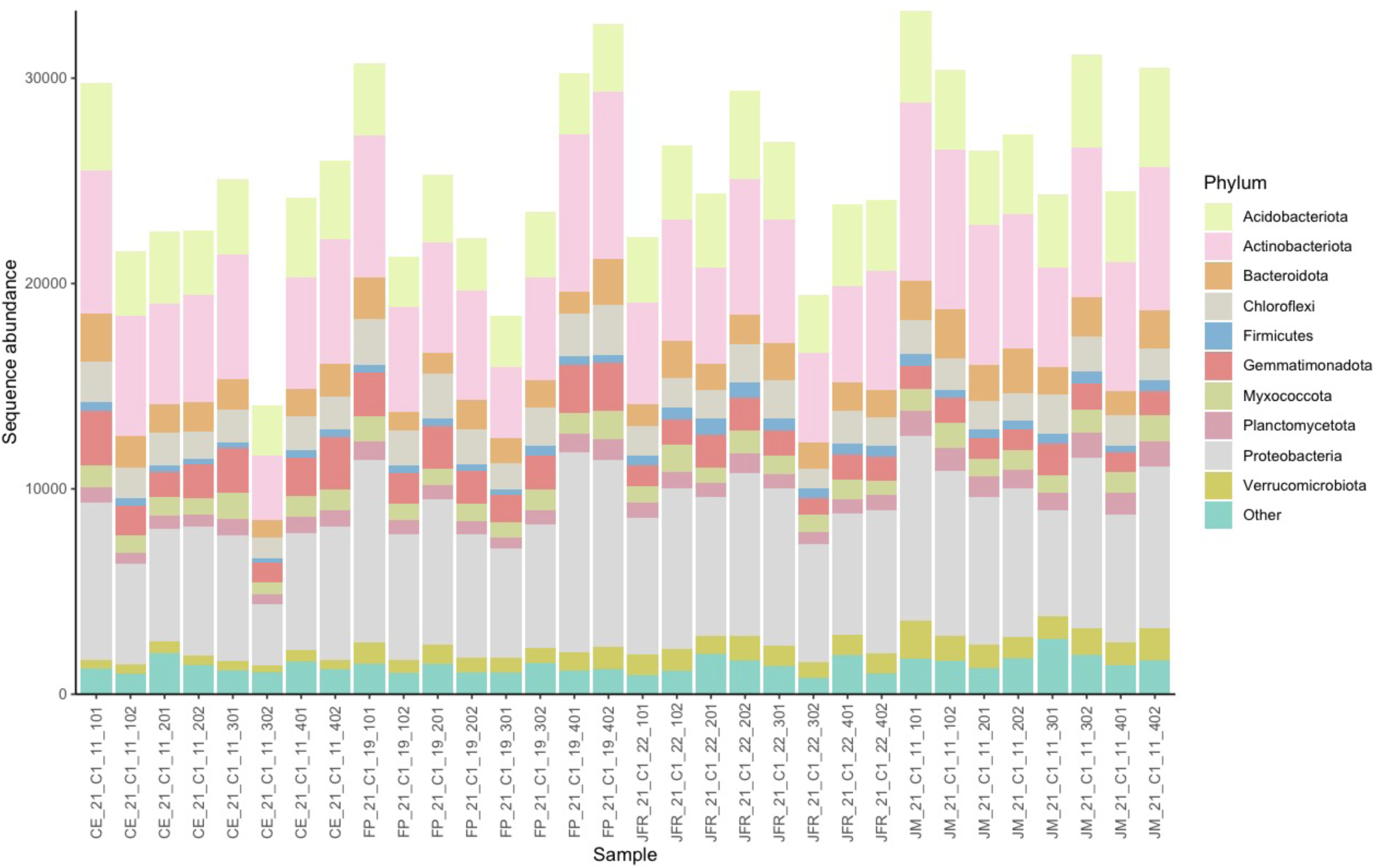
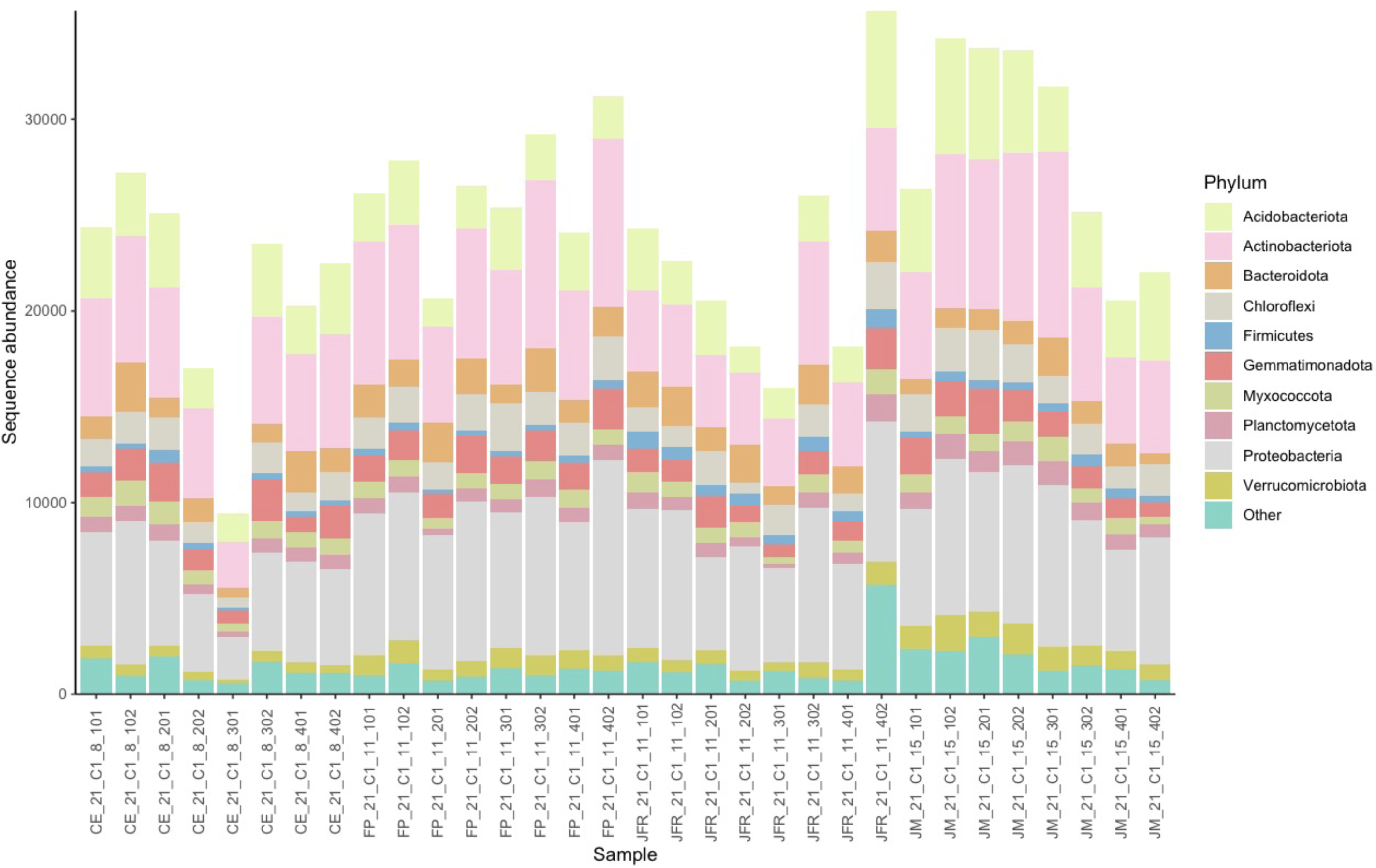
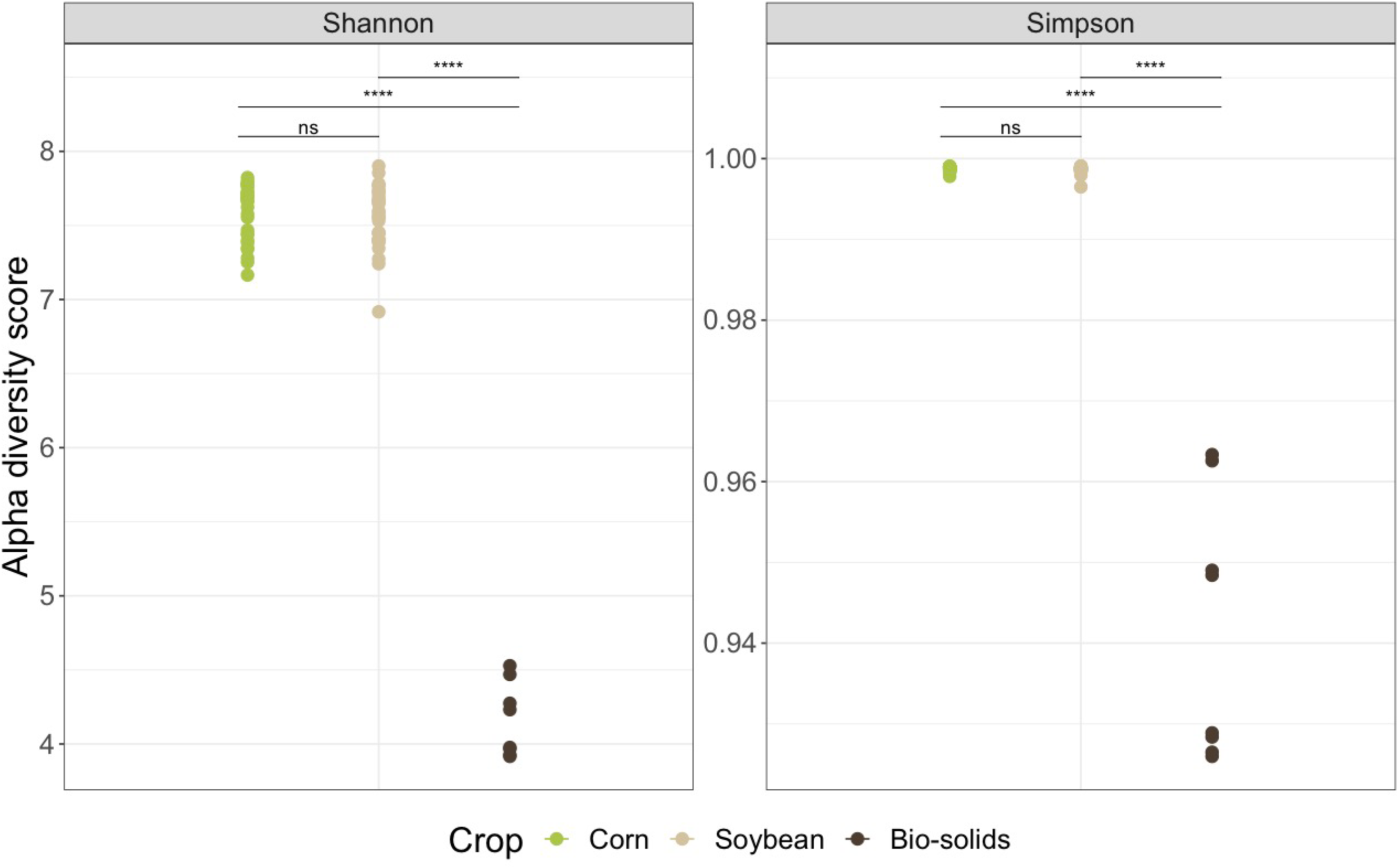
Sequence abundance of initial bacterial phyla present in each community from A) the municipal biosolids, B) corn, C) soybean and D) Shannon-Weaver and Simpson reciprocal diversity indices for soil bacterial communities. Samples were harvested during the first campaign (C1) in May 2021 from four sites in the Montérégie area of Québec. Samples are grouped by site and colour-coded by bacterial phyla in A, B, C. MBS alpha diversity was significantly lower compared to the diversity indices of corn and soybean bacterial communities for both years (Pairwise Wilcox Test *p* < 0.001). Conversely, the alpha diversity indices of corn and soybean bacterial communities were not significantly different (Pairwise Wilcox Test *p* > 0.05).

Conversely, the initial bacterial communities from the agricultural soils of corn and soybean were more complex than the MBS bacterial communities, with higher α-diversity indices (**Fig. 1D & Fig. S3**) and more ASVs (**Table S1**). In 2021, soil communities identified from the corn plots were dominated primarily by *Proteobacteria* 26.8%, *Actinobacteriota* 23.2%, and *Acidobacteriota* 13.8%, and similarly in the soil communities from soybean plots (*Proteobacteria* 26.9%, *Actinobacteriota* 23.9%, and *Acidobacteriota* 13.1%; **Fig. 1B & 1C**). In 2022, the soil communities were again similarly structured: communities from the corn plots were dominated primarily by *Actinobacteriota* 27.5%, *Proteobacteria* 26.9%, and *Acidobacteriota* 10.7%, while the communities found in soybean plots were again dominated by *Proteobacteria* 26.6%, *Actinobacteriota* 23.7% and *Acidobacteriota* 12.6% (**Fig. S2B & S2C**). Despite their high-level similarities, however, the bacterial communities found in the initial corn and soybean agricultural soils were significantly different each year (**Table 1**). We visualised this β-diversity with a Principal Component Analysis, which illustrated that the soil bacterial communities for corn and soybean were consistently off-set at each site, for both sampling years (**Fig. 2**).

**Figure 2.**
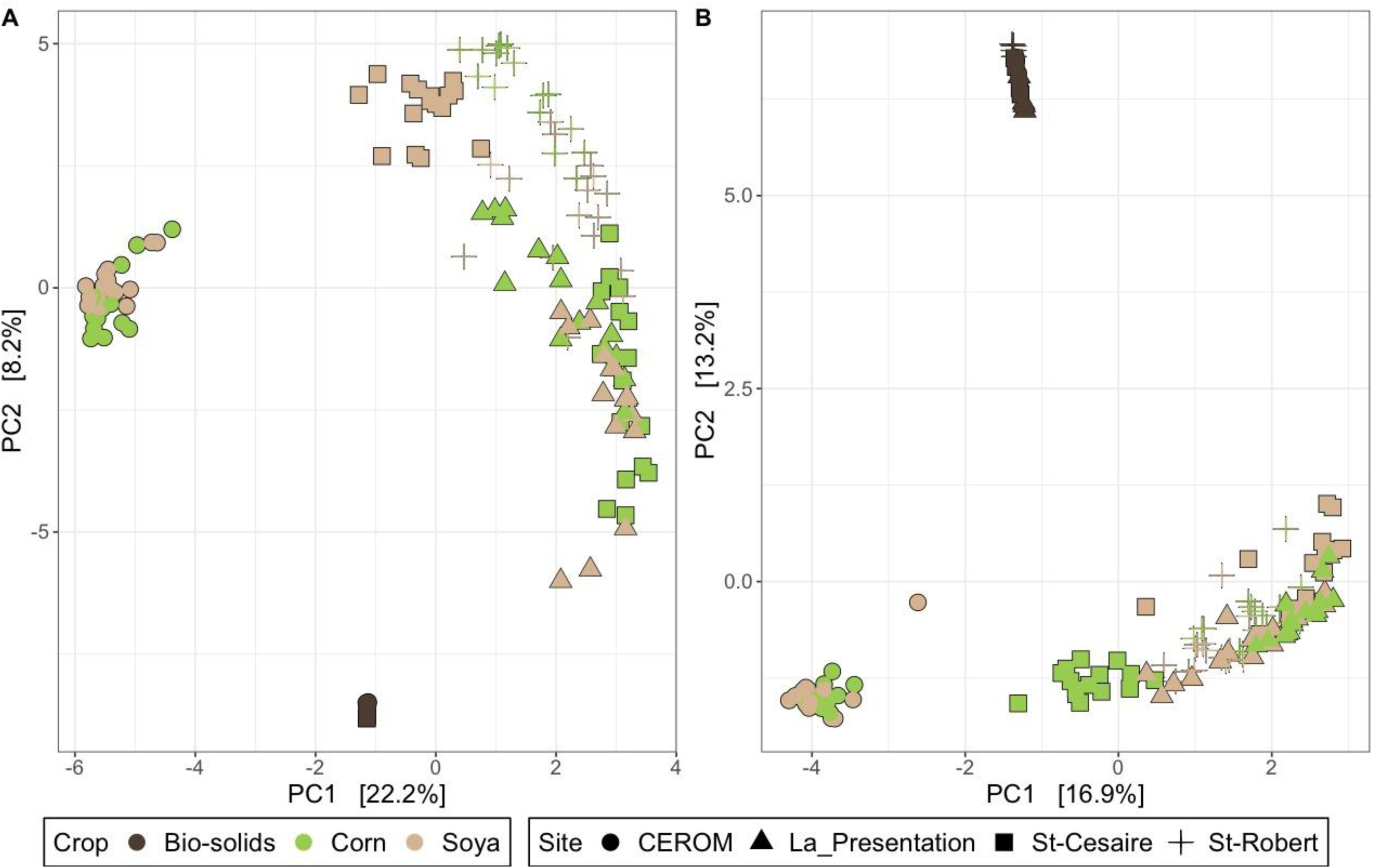
Bacterial communities between the initial municipal biosolids, and corn and soybean agricultural soils were significantly different in (A) 2021 (PERMANOVA *R^2^* = 0.078, F =8.927, *p* = 0.001), and (B) 2022 (PERMANOVA *R^2^* = 0.0140, F = 16.173, *p* = 0.001). Samples were harvested in May (C1) and July (C2) from four sites in the Montérégie area of Québec. Aitchison distances appropriate for the compositional nature of sequencing data were visualised via principal component analysis. Post-hoc analyses (pairwise PERMANOVA) indicated a near-significant difference in corn and soybean bacterial communities for 2021 (*p* = 0.082), and for 2022 (*p* = 0.076). MBS bacterial communities were consistently different from the two crop soil communities (*p* = 0.002). Samples are colour-coded according to their crop and shaped according to their site.

**Table 1.** PERMANOVA identified crops and sites as significant experimental factors for the soil bacterial communities harvested in 2021 and 2022 from four agricultural sites in the Montérégie area of Québec. PERMANOVA was calculated using an Aitchison distance matrix, with 999 permutations. We used a pairwise PERMANOVA post-hoc test to identify which crops were significantly different. The crop-site interaction was also significant.

| Experimental<br>Factors | Sample Year<sup>a</sup> |  |  |  |  |  |
| --- | --- | --- | --- | --- | --- | --- |
|  | 2021 |  |  | 2022 |  |  |
|  | F Model | R<sup>2</sup> | Pr (> F) | F Model | R<sup>2</sup> | Pr (> F) |
| Crop<sup>b</sup> | 8.927 | 0.078 | **0.001** | 16.173 | 0.140 | **0.001** |
| Site<sup>c</sup> | 21.365 | 0.280 | **0.001** | 15.124 | 0.197 | **0.001** |
| Crop ~Site | 5.271 | 0.092 | **0.001** | 3.679 | 0.096 | **0.001** |

<sup>a</sup>, Values in bold indicate significant factors or interactions
<sup>b</sup>, Bio-solids, corn or soybean
<sup>c</sup>, CEROM, La Présentation, St-Césaire, St-Robert

### 3.2 Impact of MBS treatment on corn and soybean soil bacterial communities

To test how the MBS fertiliser amendment would alter the structure of soil bacterial communities in corn and soybean agricultural sites, we compared MBS-treated soils to soils that received standard fertiliser treatments, or non-MBS treated. We found no significant differences in the α-diversity indices between soils treated with MBS, or not (**Fig. 3 & S4**), nor in terms of taxonomic composition (**Fig. S5**). We also found comparatively little overlap among ASVs identified from the initial MBS samples, and the MBS-treated and non-treated soil samples compared (**Fig. S6**). Similarly, the β-diversity analysis represented by the PCA ordination distinctly showed no difference between the MBS-treated and non-MBS-treated soil samples for both crops, in both trials, 2021 and 2022 (**Fig. 4 & S7**). This was confirmed by statistical analysis using PERMANOVA, where treatment was not significant and did not explain a high proportion of the variance in either crop, nor in either year (**Table 2 & S2**).

**Figure 3.**
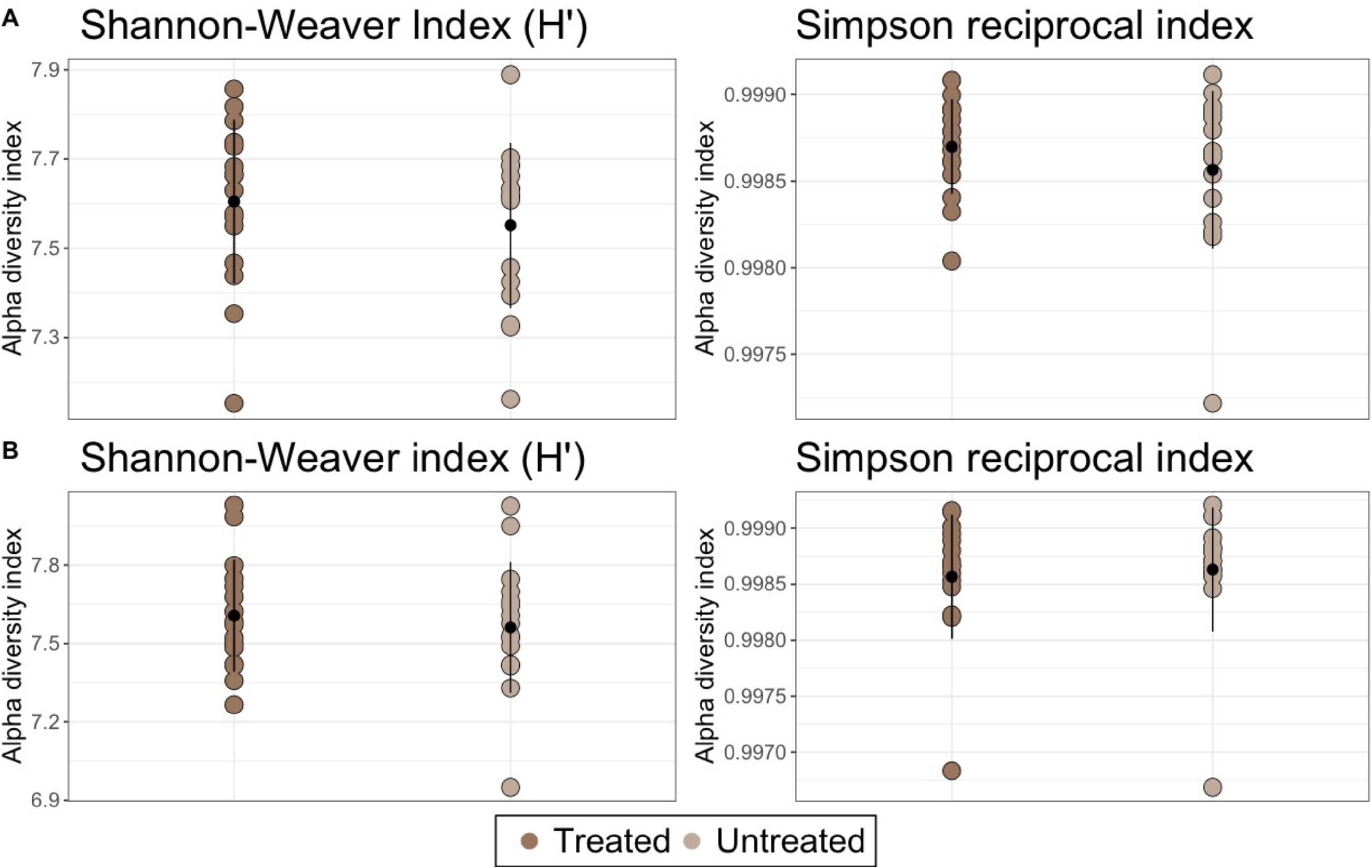
Shannon-Weaver and Simpson reciprocal diversity indices for soil bacterial communities from (A) corn and (B) soybean for the **2021** field trial were not significantly different (*p* > 0.5) between soil samples with (brown) and without MBS (green). Samples were harvested from four sites in July (C2) from the Montérégie area of Québec and bacterial composition was determined using 16S rRNA metabarcoding. A Kruskal-Wallis test was used as a statistical test.

**Figure 4.**
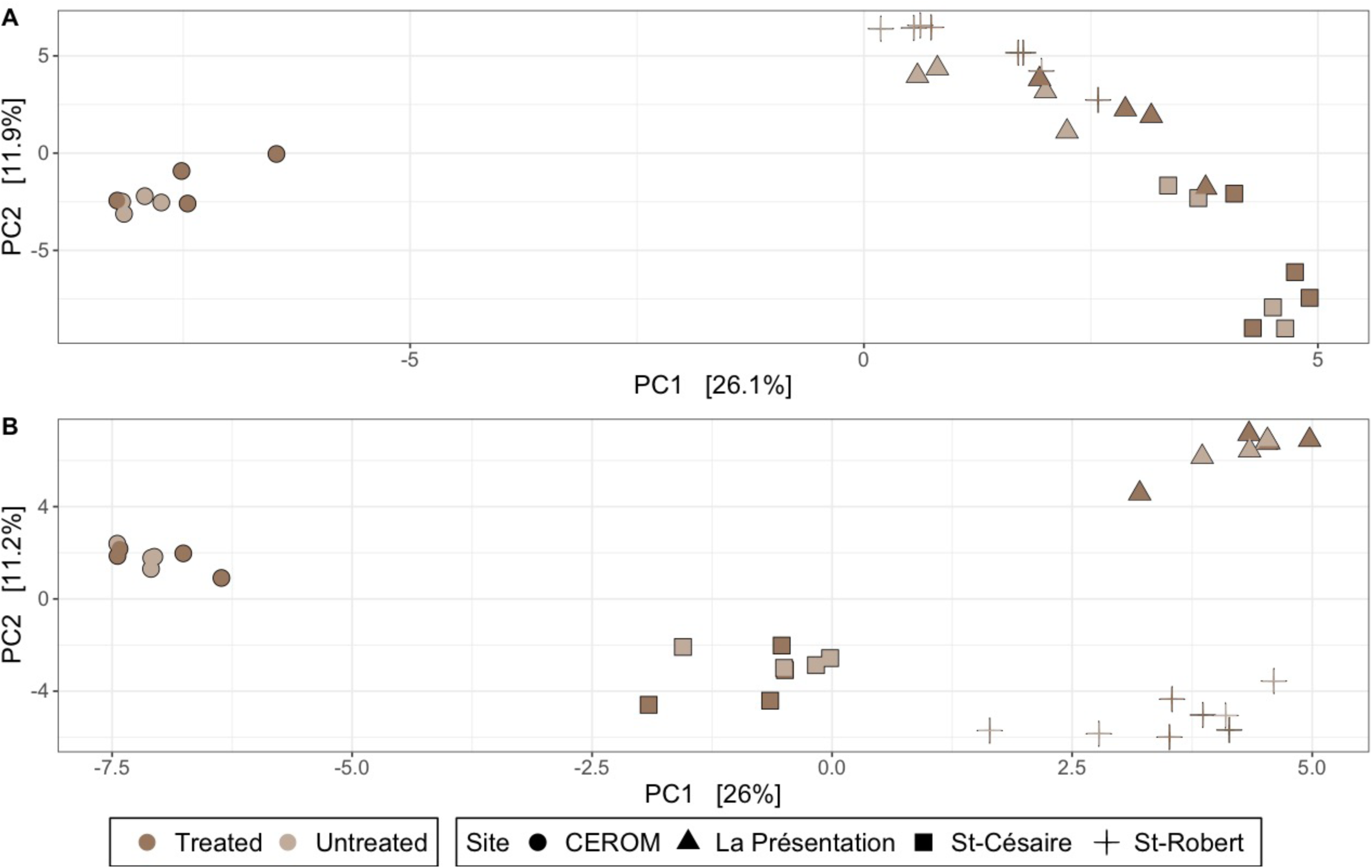
Soil bacterial communities treated with MBS from **2021** in corn (A) and soybean (B) were not significantly different from non-treated soils (PERMANOVA Corn: *R^2^* = 0.02047, F = 1.0347, *p* = 0.312; Soybean: *R^2^* = 0.02355, F = 1.0139, *p* = 0.359). Samples were harvested in July (C2) from four sites in the Montérégie area of Québec. Aitchison distances appropriate for the compositional nature of sequencing data were visualised via principal component analysis. Samples are colour-coded according to their treatments and shaped according to their site.

**Table 2.**
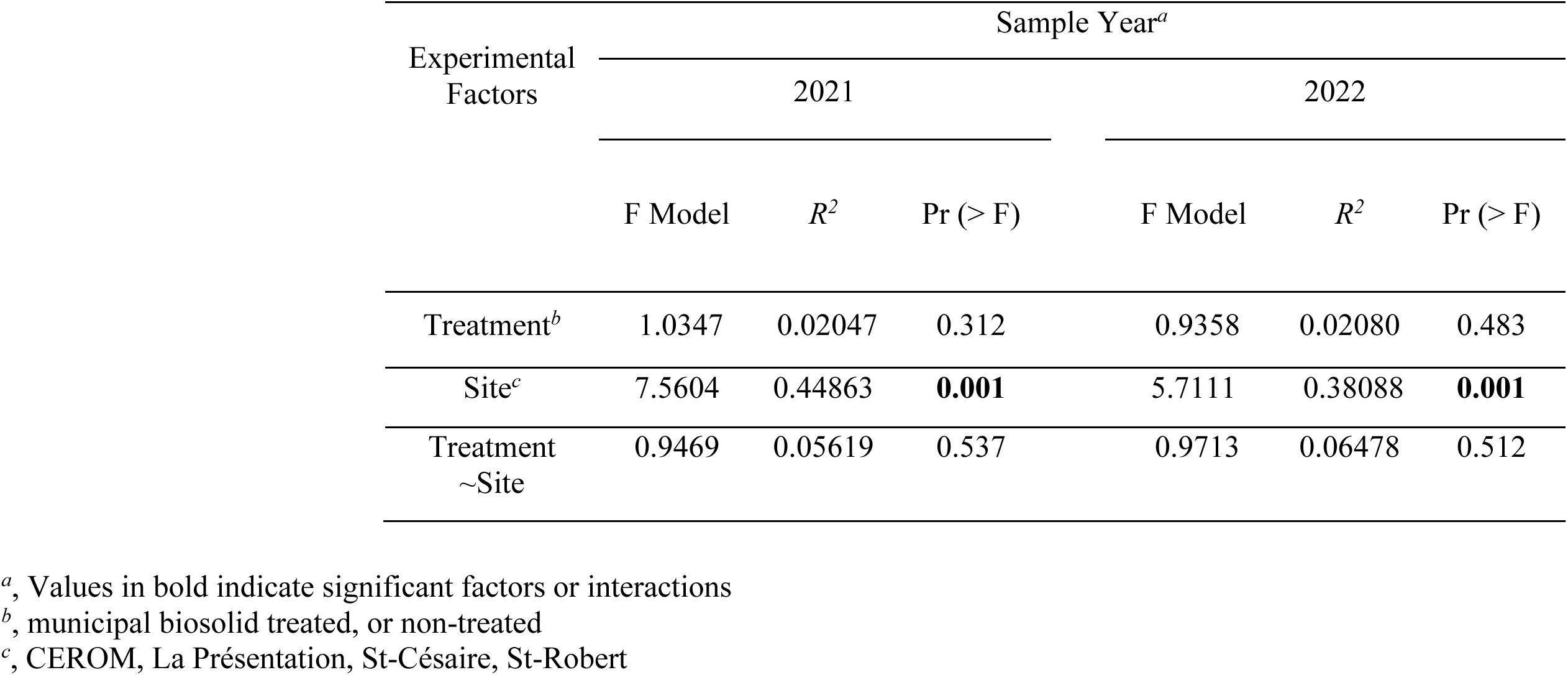
PERMANOVA identified **site** as the only significant experimental factor for the soil bacterial communities from **corn** harvested in July (C2) of 2021 and 2022 from four agricultural sites in the Montérégie area of Québec. PERMANOVA was calculated using an Aitchison distance matrix, with 999 permutations. The interaction was never significant.

### 3.3 Soil texture structurers the soil bacterial communities in corn and soybean fields

Although we found no difference among the MBS-treated and non-treated soil bacterial communities, we did observe a significant effect of the different agricultural sites on the communities throughout the experiment in both corn and soybean (**Table 2 & S2**). To further confirm the impact of the different sites on the soil bacterial communities we complemented our PERMANOVA with a distance-based redundancy analysis (db-RDA). In this analysis we quantified how the physicochemical factors of the soil--represented here by soil texture-- structured the bacterial communities at each site throughout the experiment. These models illustrated that the variation in soil textures between each site were significant in structuring the soil bacterial communities in both crops throughout the experiment (*p* = 0.001; **Fig. 5 & S8**). The RDAs explained between 19% (C2 soybean in 2022; **Fig. S8**) and 44% (C1 corn in 2021; **Fig. 5**) of the variance in the data. In corn, for example, sites containing more clay (CEROM) structured different bacterial communities from sites containing more silt (St-Cesaire and St-Robert), or sand (La Presentation; **Fig. 5**).

**Figure 5.**
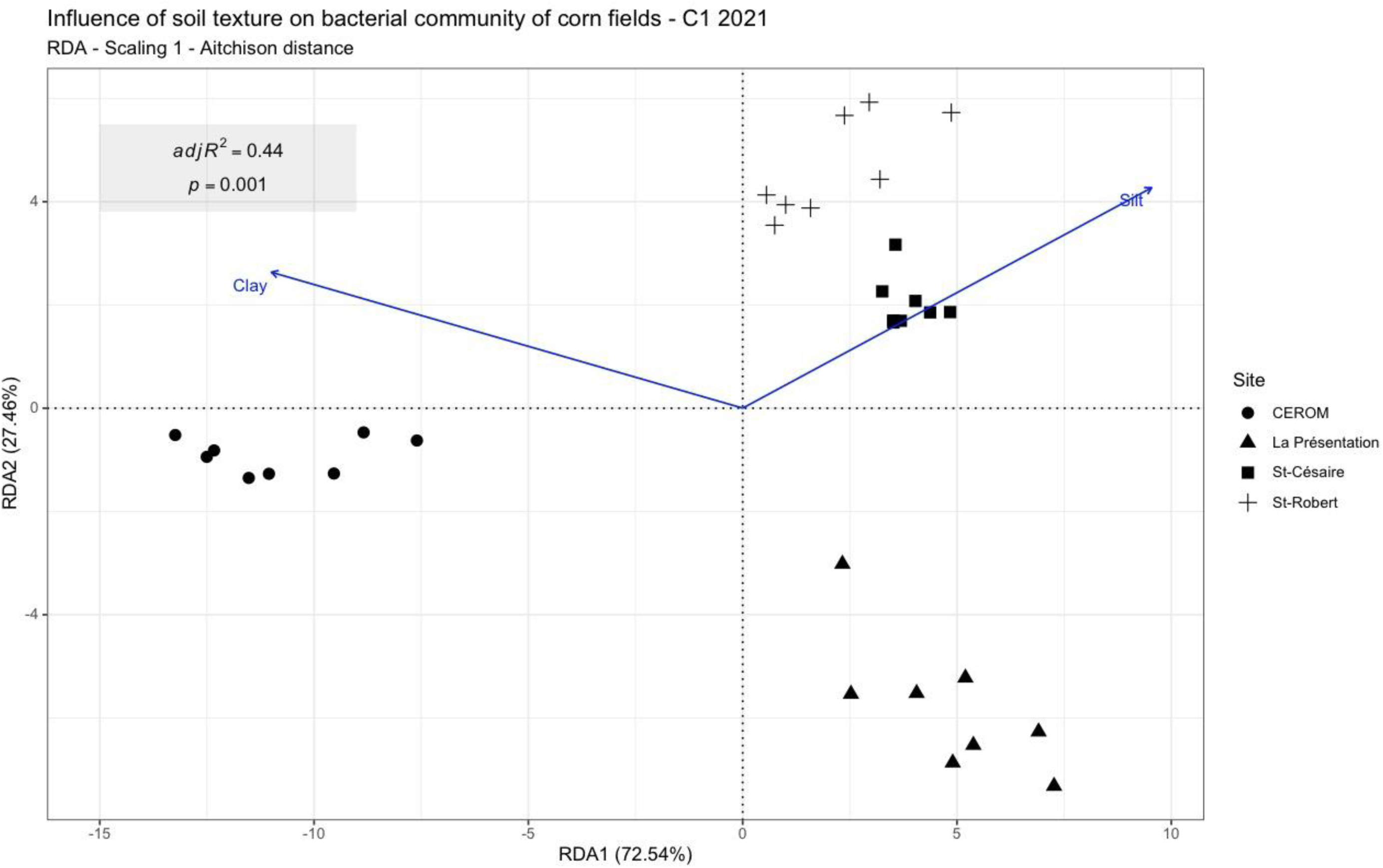
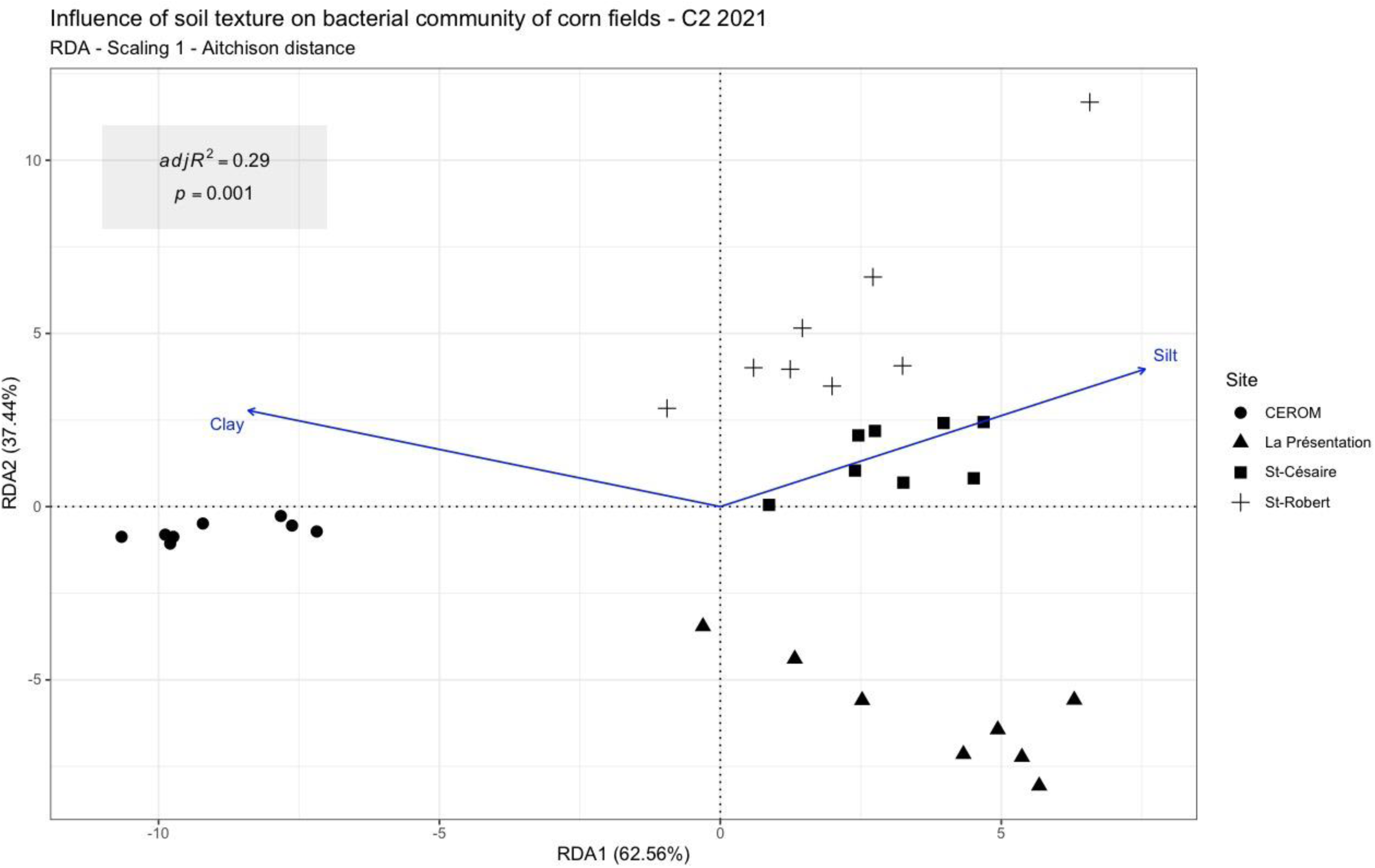
Soil bacterial communities from field experiments in **corn** plots were significantly structured by soil texture throughout 2021 (A) May, C1, (adj. *R^2^* = 0.4438, *p* = 0.001), and (B) July, C2, (adj. *R^2^* = 0.2877, *p* = 0.001). Samples were harvested at both time points from four sites in the Montérégie area of Québec. An Aitchison distance matrix, appropriate for the compositional nature of sequencing data, was used in the distance-based redundancy analysis. The RDA quantified how soil texture structured the bacterial communities, where those with similar ASV composition were plotted closer together. Soil bacterial communities from each sample are represented by different shapes for each site.

### 3.4 Detecting potential bacterial pathogenic ASVs and functions

To address the question of spreading potential human bacterial pathogens via MBS in agricultural soils, we investigated what potential pathogens or pathogenic functions could be detected in the MBS-treated soil bacterial communities. First, we found the genera *Pseudomonas*, *Stenotrophomonas,* as well as the Enterobacteriaceae family accounted for 0.7% and 2.7% of the total MBS-bacterial communities reads in 2021 and 2022 respectively (**Fig. S9**). These genera were however even less common in the agricultural soils through the experiment (**Fig. S9**).

Next, in a complementary approach, we identified potential bacterial pathogens present in the MBS and agricultural soil samples by functional annotation. First, in the MBS samples, the average proportion of ASVs with potential human pathogenic functions was low (0.45% in 2021, 0.19% in 2022), while the relative abundance of these ASVs was 0.04% in 2021 and 0.15% in 2022 (Fig. 6). The ASVs assigned to the animal parasite or symbiont functional category were also negligible (1.49% of the ASVs, 0.34% of the relative abundance in 2021, 0.47% of the ASVs and 0.55% of the relative abundance in 2022). In the agricultural soil samples, we consistently found a lower proportion both in terms of ASVs and in terms of relative abundance assigned to these functional categories. Furthermore, we found no significant increase of potential pathogenic functions when comparing the MBS-treated to the non-treated soils (**Fig. 6**).

**Figure 6.**
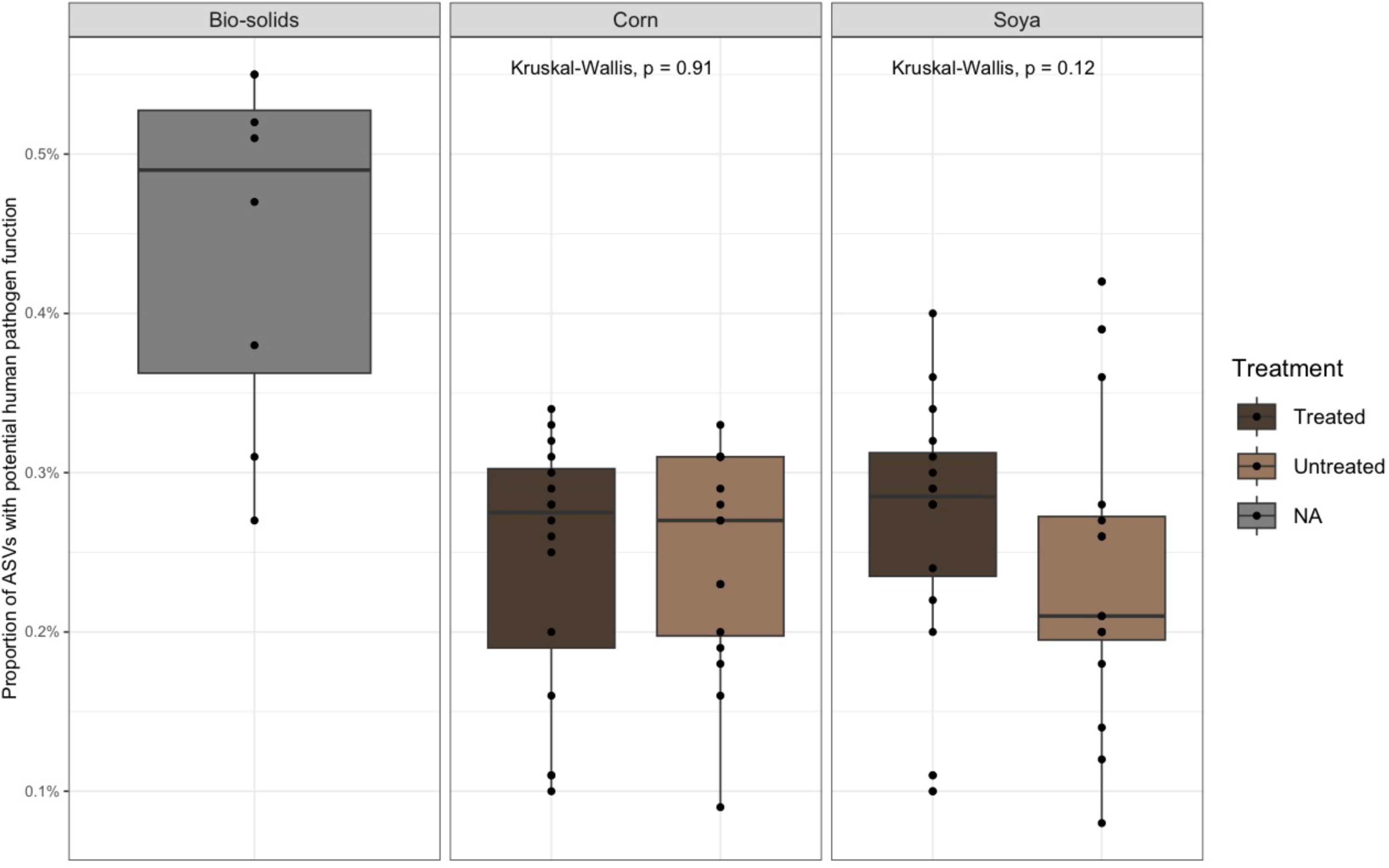
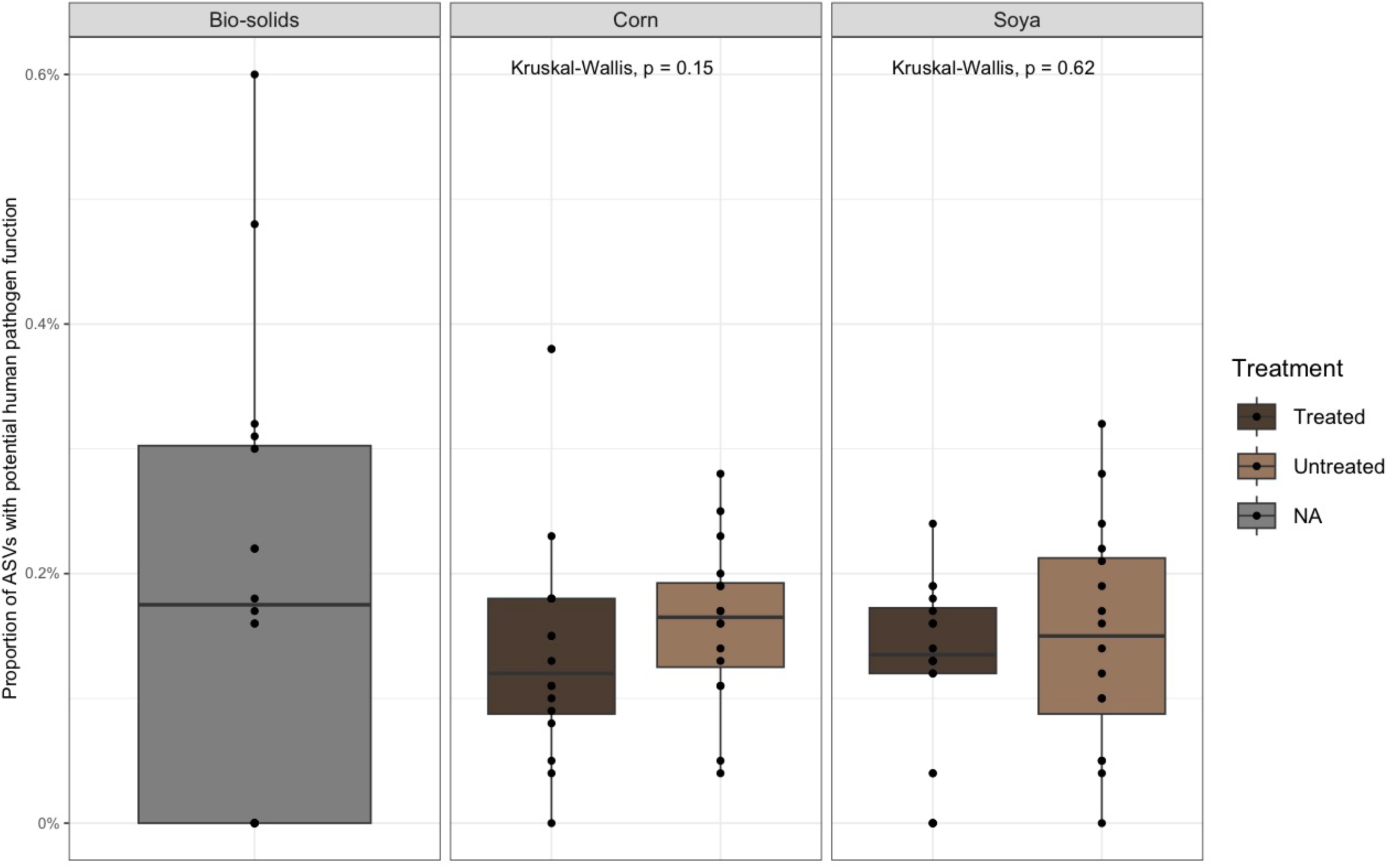
Percentage of 16S rRNA amplicon sequence variants assigned with potential human bacterial pathogenic functions in (A) 2021, and (B) 2022. Each bar is colour-coded according to sample type; the initial municipal biosolids (n = 8), the MBS-treated agricultural soils (n = 16), and the non-treated agricultural soils (n = 16). Soil samples were harvested in July (C2) from four sites in the Montérégie area of Québec. We used the non-parametric Kruskal-Wallis test to determine if there were any significant differences in the number of ASVs assigned to human pathogenic functions between MBS-treated and non-treated soils. The ASVs identified as being potential human pathogens belonged to the following genera or family: *Achromobacter, Afipia, Aquabacterium, Coxiella, Cutibacterium, Ferribacterium, Janthinobacterium, Legionella, Moraxellaceae, Niveibacterium, Oxalicibacterium, Paludibacterium, Roseomonas, Stenotrophomonas, Uliginosibacterium* and *Undibacterium*.

## 4.0 Discussion

Reducing mineral fertiliser use is a clear target for developing sustainable Canadian agriculture (Agriculture & Agri-Food Canada, 2022). Municipal biosolids (MBS) are an abundant, fertilizing residual material, rich in phosphate and nitrogen and other nutrients (Vasseur *et al.,* 1999; Warman & Termeer, 2005; Gardner *et al*., 2009; Charbonneau *et al*., 2023). However, it remains unclear how adding MBS, and its associated microbes, into agricultural systems will impact the structure and function of the resident soil bacterial communities, or potentially spread human pathogenic bacteria into the food supply-chain. In this article, we hypothesised that the use of an MBS fertiliser amendment would not significantly impact the structure nor function of the soil bacterial communities. We sampled MBS-treated and non-treated soils from corn and soybean agriculture sites and identified the composition of the soil bacterial communities via 16S rRNA metabarcoding. We found no indication that the MBS fertiliser amendment altered the structure of the soil bacterial communities. Furthermore, we identified very few ASVs associated with potential pathogenic bacteria, or potential pathogenic functions, and none belonging to the most virulent food-borne illnesses.

Our results illustrate that inoculating corn and soybean agricultural soils with MBS and their associated bacteria had no significant effect on the resident bacterial communities. We first confirmed that the initial bacterial communities identified from the MBS were significantly different from the corn and soybean soil samples (**Fig. 1, 2, S2 & S3**). We then compared MBS-treated soil bacterial communities with non-treated communities in plots of both corn and soybean. We did not find a significant effect of MBS inoculation on the α- and β-diversity indices from the soil bacterial communities (**Fig. 3, 4, S4, & S6**). This could suggest that our MBS-treatment was simply insufficient, or perhaps not concentrated enough, or that the total amount of new bacteria added to the soil was small compared to the original community, in order to cause any change to the structure of the soil bacterial communities. In this instance, however, a weak treatment is not a deficit in our study. Here, our goal was to determine the impact of MBS as a fertiliser amendment on soil bacterial communities in agricultural systems. The work of Charbonneau *et al*., (2023) clearly illustrated that the MBS-treatment was functionally sufficient as a fertiliser amendment; both MBS-treated and non-treated plots of corn and soybean produced similar yields. Although higher doses of MBS may have evoked obvious shifts among the soil bacterial communities, they would be unrealistic of the amount of MBS permitted to be applied in agricultural systems, as far as phosphorus amendment is concerned (Parent & Gagné, 2010).

Moreover, that the MBS-associated bacteria were not more successful in colonising the agricultural soils speaks to the resilience of the resident soil bacterial communities (Hong *et al*., 2021; Yates *et al*., 2022). The bacterial pathogens responsible for most food-borne illnesses, enterohemorrhagic *Escherichia coli*, *Salmonella enterica* and *Listeria monocytogenes*, all have large genomes, are metabolically diverse, and are well-adapted to grow and compete in soil environments (Brennan *et al*., 2022). The same is also true for many soil-born, opportunistic bacterial pathogens, such as *Pseudomonas aeruginosa*, or *Stenotrophomonas maltophilia* (Rossi *et al*., 2021; Brooke, 2021). Furthermore, soils are well-known reservoirs for potential human bacterial pathogens and pathogenic functions, such as virulent gene cassettes (Scott *et al*., 2022). Therefore, it would be reasonable to find some potential human bacterial pathogens lurking in the agricultural soils we sampled irrespective of the applied MBS.

As we illustrated, the bacterial communities from various agricultural sites did contain a small fraction of known potential bacterial pathogens; consistently less than 0.7% of the ASVs (**Fig. 6 & S9**). However, given that the MBS-treated and non-treated agricultural soils remained indistinct, it is likely that these few potential bacterial pathogens belong to the extant soil reservoir of background, opportunistic pathogens that would be found in any soil (Brennan *et al*., 2022; Banerjee & van der Heijden, 2023). This suggests that little of the MBS bacterial community was actually able to find an exploitable niche in the soil communities and establish resident populations (Yates *et al*., 2022). Perhaps with long-term usage, the impact would accrue.

Overall, the abundant taxa found in MBS samples are typical of municipal wastewater with *Bacteriodota*, *Firmicutes, Actinobacteriota*, and *Proteobacteria*, as the most abundant phyla (Numberger *et al.,* 2019; LaMartina *et al*., 2021; Brison *et al*. 2022). Furthermore, that we detected very few known, zoonotic bacterial pathogens in general throughout the experiment could be attributed to how the MBS are prepared. The current policy in Québec requires that before batches of MBS can be taken, they must pass a coliform test. As such, there ought to have been a minimal amount of potential human pathogens present at the outset. Therefore, the MBS should pose no more a biosecurity threat than public swimming areas, which are similarly monitored.

## 5.0 Conclusion

Municipal biosolids (MBS) are effective in reducing mineral fertiliser use, since they are an abundant residual organic material, rich in phosphate and nitrogen and other oligo-nutrients, as shown for various crops. Here, we addressed two out-standing questions in using MBS in agricultural systems: do MBS, and its associated microbes, impact the structure and function of the resident soil bacterial communities? Does applying MBS in agricultural fields spread potential pathogenic bacteria into the food supply-chain? We tested the hypothesis that the use of an MBS fertiliser amendment would not significantly impact the structure nor function of the soil bacterial communities. Using 16S rRNA metabarcoding to identify the bacteria present, we found no indication that the MBS fertiliser amendment altered the structure of the soil bacterial communities after two years of monitoring. Furthermore, we identified very few ASVs associated with potential pathogenic bacteria, or potential pathogenic functions. Rather, our data suggests that MBS bacterial communities do not persist in the soil communities, and that different soils continue to be more significant factors in structuring bacterial soil communities than any MBS inoculation. Therefore, from a community composition, structure, and function point-of-view, using MBS in agriculture poses no greater concern to the public than existing soil bacterial communities.

## Supporting information

Blakney et al., 2023 BioRxiv Supplemental Materials

## Author Contributions

All authors contributed to the study conception and design. Funding acquisition as well as the provision of resources were provided by M.L. Material preparation, soil sampling, and data collection were performed by A.C., and M.M. Sequence analysis was done by E.G. Statistical analyses, figure preparation, and the article were prepared by S.M. and A.B. All authors commented on previous versions of the paper. All authors have read and agreed to the published version of the paper.

## Informed Consent Statement

Informed consent was obtained from all subjects involved in the study.

## Data Availability Statement

The data presented in this study are available within the article or on request from Marc Lucotte. Sequencing data and metadata are available at NCBI Bioproject under accession number: PRJNA1044782.

## Footnotes

1 Abbreviations: municipal biosolids, MBS

## References

Agriculture & Agri-Food Canada (2022) Fertilizer emissions reduction target. https://agriculture.canada.ca/en/department/transparency/public-opinion-research-consultations/share-ideas-fertilizer-emissions-reduction-target

Banerjee S, & van der Heijden MGA (2023) Soil microbiomes and one health. Nature Reviews Microbiology 21:6–20.

Bell TH, Stefani FOP, Abram K, Champagne J, Yergeau É, Hijri M, & St-Arnaud M (2016) A diverse soil microbiome degrades more crude oil than specialized bacterial assemblages obtained in culture. Appl Environ Microbiol 82(18): 5530–5541.

Blakney AJC, Bainard LD, St-Arnaud M, & Hijri M (2022) *Brassicaceae* host plants mask the feedback from the previous year’s soil history on bacterial communities, except when they experience drought. Environmental Microbiology 24:3529–3548.

Brennan FP, Alsanius BW, Allende A, Burgess CM, Moreira H, Johannessen GS, Castro PML, Uyttendaele M, Truchado P, & Holden NJ (2022) Harnessing agricultural microbiomes for human pathogen control. ISME Communications 2:44.

Brison A, Rossi P, Gelb A, & Derlon N (2022) The capture technology matters: composition of municipal wastewater solids drives complexity of microbial community structure and volatile fatty acid profile during anaerobic fermentation. Science of the Total Environment, 815, 152762.

Brooke JS (2021) Advances in the Microbiology of *Stenotrophomonas maltophilia*. Clinical Microbiology Reviews 34:e00030–19.

Callahan BJ, McMurdie PJ, Rosen MJ, Han AW, Johnson AJ, & Holmes SP (2016) DADA2: High-resolution sample inference from Illumina amplicon data. Nature Methods 13:581–583.

Charbonneau A, Lucotte M, Moingt M, Blakney AJC, Morvan S, Bipfubusa M, & Pitre FE (*Submitted*) Fertilisation of agricultural soils with municipal biosolids: Part 1: glyphosate and aminomethylphosphonic acid inputs on Québec field crop soils.

Crippa M, Solazzo E, Guizzardi D, Monforti-Ferrario F, Tubiello FN, & Leip A (2021) Food systems are responsible for a third of global anthropogenic GHG emissions. Nature Food 2:198–209.

Depoorter E, Bull MJ, Peeters C, Coenye T, Vandamme P, & Mahenthiralingam E (2016) *Burkholderia*: an update on taxonomy and biotechnological potential as antibiotic producers. Applied Microbiology & Biotechnology 100:5215–5229.

Fargione JE, Bassett S, Boucher T et al., (2018) Natural climate solutions for the United States. Science Advances 4:eaat1869.

Gloor GB, Macklaim JM, Pawlowsky-Glahn V, & Egozcue JJ (2017). Microbiome datasets are compositional: and this is not optional. Frontiers in Microbiology 8:2224.

Halpern BS, Frazier M, Verstaen J, et al (2022) The environmental footprint of global food production. Nature Sustainability 5:1027–1039.

Hong P, Schmid B, De Laender F, Eisenhauer N, Zhang X, Chen H, Craven D, De Boeck HJ, Hautier Y, Petchey OL, Reich PB, Steudel B, Striebel M, Thakur MP, & Wang S (2021) Biodiversity promotes ecosystem functioning despite environmental change. Ecology Letters 25:555 –569.

Hou S, Thiergart T, Vannier N, Mesny F, Ziegler J, Pickel B, & Hacquard S (2021) A microbiota-root–shoot circuit favours *Arabidopsis* growth over defence under suboptimal light. Nature Plants 7:1078–1092.

Keeler BL, Gourevitch JD, Polasky S, Isbell F, Tessum CW, Hill JD, & Marshall JD (2016) The social cost of nitrogen. Science Advances 2:e1600219.

Klindworth A, Pruesse E, Schweer T, Peplies J, Quast C, Horn M, & Glöckner FO (2012) Evaluation of general 16S ribosomal RNA gene PCR primers for classical and next-generation sequencing-based diversity studies. Nucleic Acids Research 41:1–11.

LaMartina EL, Mohaimani AA, & Newton RJ (2021) Urban wastewater bacterial communities assemble into seasonal steady states. Microbiome 9:116.

Lay CY, Bell TH, Hamel C, Harker KN, Mohr R, Greer CW, Yergeau É, & St-Arnaud M (2018) Canola root–associated microbiomes in the Canadian prairies. Frontiers in Microbiology 9:1–19.

Lau JA, & Lennon JT (2012) Rapid responses of soil microorganisms improve plant fitness in novel environments. Proceedings of the National Academy of Sciences of the United States of America 109:14058–14062.

Liu C, Cui Y, Li X, & Yao M (2021) *microeco*: an R package for data mining in microbial community ecology. FEMS Microbiology Ecology 97:fiaa255.

Louca S, Parfrey LW, & Doebeli M (2016). Decoupling function and taxonomy in the global ocean microbiome. Science, 353(6305), 1272–1277.

Marasco R, Rolli E, Ettoumi B, Vigani G, Mapelli F, Borin S, Abou-Hadid AF, El-Behairy UA, Sorlini C, Cherif A, Zocchi G, & Daffonchio D (2012) A drought resistance-promoting microbiome is selected by root system under desert farming. PLOS One 7:e48479.

Martin M (2011) Cutadapt removes adapter sequences from high-throughput sequencing reads. EMBnet.Journal 17:10–12.

McMurdie P, & Holmes S (2013) Phyloseq: an R package for reproducible interactive analysis and graphics of microbiome census data. PLOS One 8:e61217.

Mendes R, Kruijt M, de Bruijn I, Dekkers E, van der Voort M, Schneider JHM, Piceno YM, DeSantis TZ, Andersen GL, Bakker PAHM, & Raaijmaker JM (2011) Deciphering the rhizosphere microbiome for disease-suppressive bacteria. Science 332:1097–1100.

Milner AM, & Boyd IL (2017) Toward pesticidovigilance. Science 357:1232–1234.

Numberger D, Ganzert L, Zoccarato L, Mühldorfer K, Sauer S, Grossart HP, & Greenwood AD (2019) Characterization of bacterial communities in wastewater with enhanced taxonomic resolution by full-length 16S rRNA sequencing. Scientific reports 9:9673.

Oksanen J, Blanchet FG, Friendly M, Kindt R, Legendre P, McGlinn D, Minchin PR, O’Hara RB, Simpson GL, Solymos P, Stevens MHH, Szoecs E, & Wagner H (2020) Vegan: community ecology package. R package version 2.5–7.

Parent LE, & Gagné G, éditeurs scientifiques (2010) Guide de référence en fertilisation, 2e édition. Centre de référence en agriculture et agroalimentaire du Québec, Québec.

Parks DH, Chuvochina M, Waite DW, Rinke C, Skarshewski A, Chaumeil PA, & Hugenholtz P (2018) A standardized bacterial taxonomy based on genome phylogeny substantially revises the tree of life. Nature Biotechnology 36:996–1004.

Quinn TP, Erb I, Richardson MF, & Crowley TM (2018) Understanding sequencing data as compositions: an outlook and review. Bioinformatics 34:2870–2878.

R Core Team (2020). R: A language and environment for statistical computing. R Foundation for Statistical Computing, Vienna, Austria.

Richardson AE, Barea JM, McNeill AM, & Prigent-Combaret C (2009) Acquisition of phosphorus and nitrogen in the rhizosphere and plant growth promotion by microorganisms. Plant & Soil 321:305–339.

Rosegrant MW, Ringler C, & Zhu T (2009) Water for agriculture: maintaining food security under growing scarcity. Annual Review of Environment and Resources 34:205–222.

Rossi E, La Rosa R, Bartell JA, Marvig RL, Haagensen JAJ, Sommer LM, Molin S, & Johansen HK (2021) *Pseudomonas aeruginosa* adaptation and evolution in patients with cystic fibrosis. Nature Reviews Microbiology 19:331–342.

Roy S, Coldren C, Karunamurthy A, Kip NS, Klee EW, Lincoln SE, Leon A, Pullambhatla M, Temple-Smolkin RL, Voelkerding KV, Wang C, & Carter AB (2018) Standards and guidelines for validating next-generation sequencing bioinformatics pipelines. Journal of Molecular Diagnostics 20:4–27.

Ryan RP, Monchy S, Cardinale M, Taghavi S, Crossman L, Avison MB, Berg G, van der Lelie D, & Dow JM (2009) The versatility and adaptation of bacteria from the genus *Stenotrophomonas*. Nature Review Microbiology 7:514–525.

Schlatter DC, Paul NC, Shah DH, Shah DH, Schillinger WF, Bary AI, Sharratt B, & Paulitz TC (2019) Biosolids and tillage practices influence soil bacterial communities in dryland wheat. Microbial Ecology 78:737–752.

Scott A, Murray R, Tien YC, & Topp E (2022) Contamination of hay and haylage with enteric bacteria and selected antibiotic resistance genes following fertilization with dairy manure or biosolids. Canadian Journal of Microbiology 68:249–257.

Sikes BA, Cottenie K, & Klironomos JN (2009) Plant and fungal identity determines pathogen protection of plant roots by arbuscular mycorrhizas. Journal of Ecology 97:1274–1280.

Strickland MS, Osburn E, Lauber C, Fierer N, & Bradford MA (2009) Litter quality is in the eye of the beholder: initial decomposition rates as a function of inoculum characteristics. Functional Ecology 23:627 –636.

Tubiello FN, Salvatore M, Ferrara AF, et al (2015) The contribution of agriculture, forestry and other land use activities to global warming, 1990-2012. Global Change Biology 21:2655–2660.

Walsh CM, Becker-Uncapher I, Carlson M, & Fierer N (2021) Variable influences of soil and seed-associated bacterial communities on the assembly of seedling microbiomes. ISME J 15:2748–2762.

Walterson AM, & Stavrinides J (2015) *Pantoea*: insights into a highly versatile and diverse genus within the *Enterobacteriaceae*. FEMS Microbiology Review 39:968–984.

Weidner S, Koller R, Latz E, Kowalchuk G, Bonkowski M, Scheu S, & Jousset A (2015) Bacterial diversity amplifies nutrient-based plant–soil feedbacks. Functional Ecology 29:1341–1349.

Wickham H (2016) ggplot2: elegant graphics for data analysis. Springer-Verlag New York.

Wittwer RA, Bender SF, Hartman K, Hydbom S, Lima RAA, Loaiza V, Nemecek T, Oehl F, Olsson PA, Petchey O, Prechsl UE, Schlaeppi K, Scholten T, Seitz S, Six J, & van der Heijden MGA (2021) Organic and conservation agriculture promote ecosystem multifunctionality. Science Advances 7:eabg6995.

Yates CF, Trexler RV, Bonet I, King WL, Hockett KL, & Bell TH (2022) Rapid niche shifts in bacteria following conditioning in novel soil environments. Functional Ecology 00:1–11

Yilmaz P, Parfrey LW, Yarza P, Gerken J, Pruesse E, Quast C, Schweer T, Peplies J, Ludwig W, & Glöckner FO (2013) The SILVA and “all-species living tree project (LTP)” taxonomic frameworks. Nucleic Acids Research 42:D643–D648.

Yu P, He X, Baer M, Beirinckx S, Tian T, Moya YAT, Zhang X, Deichmann M, Frey FP, Bresgen V, Li C, Razavi BS, Schaaf G, von Wirén N, Su Z, Bucher M, Tsuda K, Goormachtig S, Chen X, & Hochholdinger F (2021) Plant flavones enrich rhizosphere *Oxalobacteraceae* to improve maize performance under nitrogen deprivation. Nature Plants 7:481–499.

Zou T, Zhang X, & Davidson EA (2022) Global trends of cropland phosphorus use and sustainability challenges. Nature 611:81–87.

