## Supplementary material for "Fertilisation of agricultural soils with municipal biosoilds: Part 2- Site properties, environmental factors, and crop identify influence soil bacterial communities more than municipal biosolid application": Blakney et al., 2023 BioRxiv Supplemental Materials

**Table S1**. The soil bacterial communities consistently yielded more unique ASVs than the municipal biosolid samples (MBS) over both years of the experiment, regardless of crop (corn or soybean), or sampling date (C1, May; C2, July). Samples were harvested throughout the 2021 and 2022 growing seasons from four different agricultural sites in the Montérégie area of southern Québec. We obtained the raw 16S rRNA reads from the Illumina MiSeq platform at Génome Québec and processed them through DADA2. We retained 3 481 870 reads for 2021 and 773 403 reads for 2022 (16S rRNA reads reported here) for ASV inference. A total of 34 142 and 7 886 ASVs were identified in the 2021 and 2022 datasets, respectively.

| | | 2021 | |
|---|---|---|---|
| | Sampling Date[a] | Total number of 16S rRNA reads (mean number of reads per sample)[b] | ASV Occurrence |
| MBS | / | 248 576 (31072 ± 4566) | 1184 |
| Corn | C1 | 814 767 (25461 ± 4351) | 27 750 |
| | C2 | 826 386 (25825 ± 6469) | 27 753 |
| Soybean | C1 | 789 437 (24670 ± 5828) | 27 877 |
| | C2 | 802 704 (25085 ± 4539) | 28 013 |
| | | Total Reads = 3 481 870 | Total Unique = 34 142 |

| | | 2022 | |
|---|---|---|---|
| | Sampling Date[a] | Total number of 16S rRNA reads (mean number of reads per sample)[b] | ASV Occurrence |
| MBS | / | 89 404 (5587 ± 476) | 983 |
| Corn | C1 | 170 535 (5501 ± 846) | 7113 |
| | C2 | 165 374 (5167 ± 835) | 7095 |
| Soybean | C1 | 176 066 (5502 ± 948) | 7075 |
| | C2 | 172 024 (5376 ± 814) | 7132 |
| | | Total Reads = 773 403 | Total Unique = 7886 |

[a], C1, May; C2, July

[b], Values are presented as mean ± SD

**Table S2**. PERMANOVA identified **site** as the only significant experimental factor for the soil bacterial communities from **soybean** harvested in July (C2) of 2021 and 2022 from four agricultural sites in the Montérégie area of Québec. PERMANOVA was calculated using an Aitchison distance matrix, with 999 permutations. The interaction was never significant.

| Experimental Factors | Sample Year[a] | | | | | |
|---|---|---|---|---|---|---|
| | 2021 | | | 2022 | | |
| | F Model | $R^2$ | Pr (> F) | F Model | $R^2$ | Pr (> F) |
| Treatment[b] | 0.9252 | 0.01842 | 0.457 | 1.0139 | 0.02355 | 0.359 |
| Site[c] | 7.5195 | 0.44912 | **0.001** | 5.0192 | 0.34969 | **0.001** |
| Treatment ~Site | 0.9148 | 0.05464 | 0.571 | 0.9961 | 0.06940 | 0.443 |

[a], Values in bold indicate significant factors or interactions
[b], municipal biosolid treated, or non-treated
[c], CEROM, La Présentation, St-Césaire, St-Robert

**Figure S1** A)

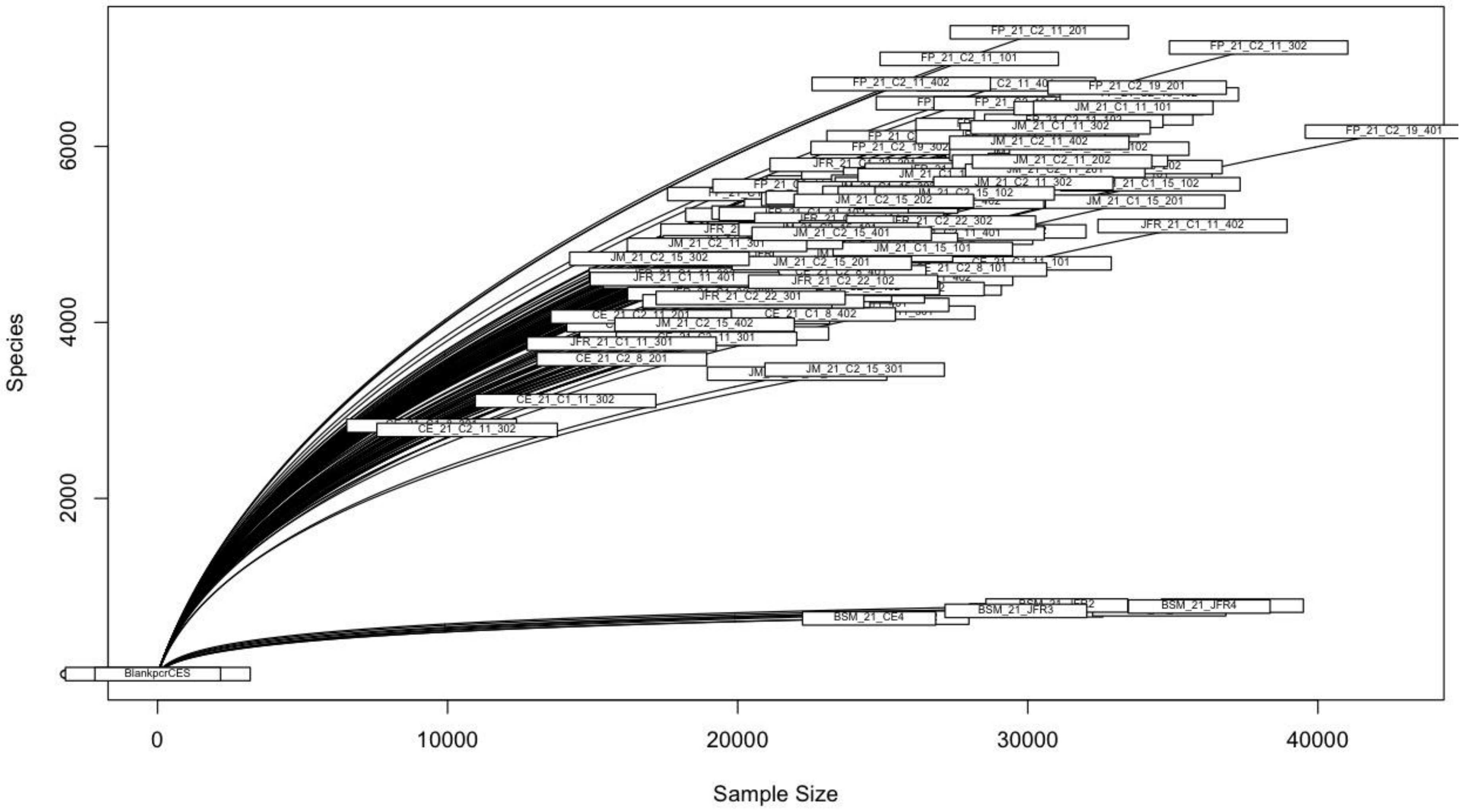
Species
Sample Size
0
2000
4000
6000
0
10000
20000
30000
40000
FP_21_C2_11_201
FP_21_C2_11_302
FP_21_C2_11_101
FP_21_C2_11_402
FP_21_C2_19_201
JM_21_C1_11_101
FP_21_C2_19_401
JM_21_C1_11_302
JM_21_C2_11_402
FP_21_C2_19_302
JM_21_C2_11_202
JM_21_C2_11_302
JM_21_C2_15_202
JM_21_C1_15_201
JFR_21_C1_11_402
JM_21_C2_15_401
JM_21_C2_11_301
JM_21_C1_15_101
JM_21_C2_15_302
JM_21_C2_15_201
JFR_21_C1_11_401
JFR_21_C2_22_102
JFR_21_C2_22_301
CE_21_C1_8_402
JM_21_C2_15_402
JFR_21_C1_11_301
CE_21_C2_8_201
JM_21_C2_15_301
CE_21_C1_11_302
CE_21_C2_11_302
BSM_21_JFR4
BSM_21_JFR3
BSM_21_CE4
BlankpcrCES

**Figure S1** B)

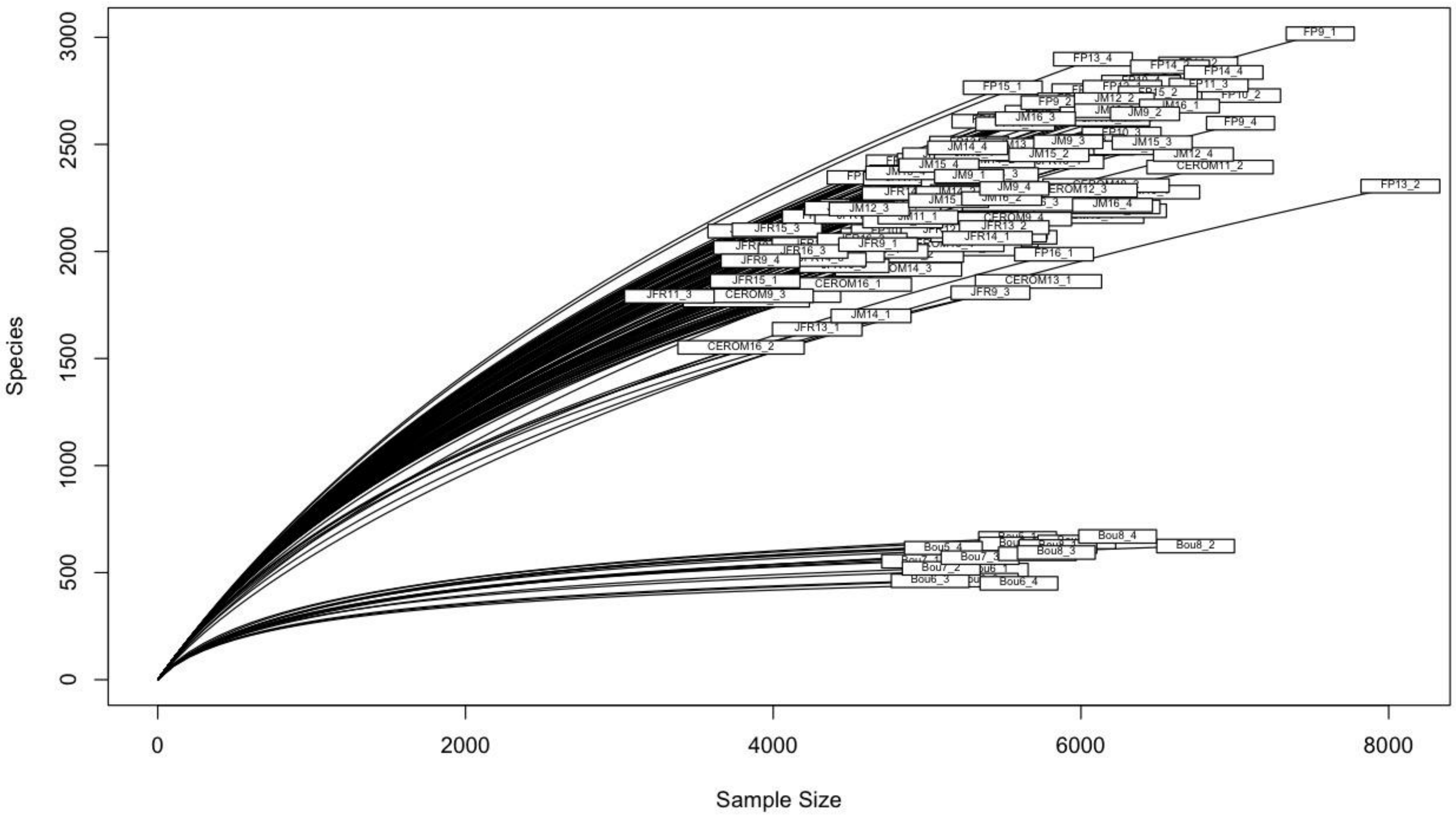
Species
Sample Size
0
500
1000
1500
2000
2500
3000
0
2000
4000
6000
8000
FP9_1
FP13_4
FP14_4
FP15_1
FP9_2
FP11_3
FP10_2
JM16_3
JM9_2
JM16_1
FP9_4
JM14_4
JM9_3
JM15_3
JM15_2
JM12_4
CEROM11_2
FP13_2
JM15_4
JM9_1
JM9_4
JM12_3
JM16_4
JM11_1
JFR15_3
JFR13_2
JFR14_1
JFR9_1
JFR16_3
JFR9_4
FP16_1
JFR15_1
CEROM16_1
CEROM13_1
JFR11_3
CEROM9_3
JFR9_3
JM14_1
JFR13_1
CEROM16_2
Bou8_4
Bou8_2
Bou5_4
Bou7_3
Bou8_3
Bou7_2
Bou6_3
Bou6_4

**Fig S1**. Rarefaction curves illustrated that the majority of the bacterial communities were identified in the municipal biosolids from both 2021 (A) and 2022 (B), while nearly all the communities were captured from the agricultural soil samples harvested in both years, from four different sites around the Montérégie area of Québec.

**Figure S2** A) 2022 MBS

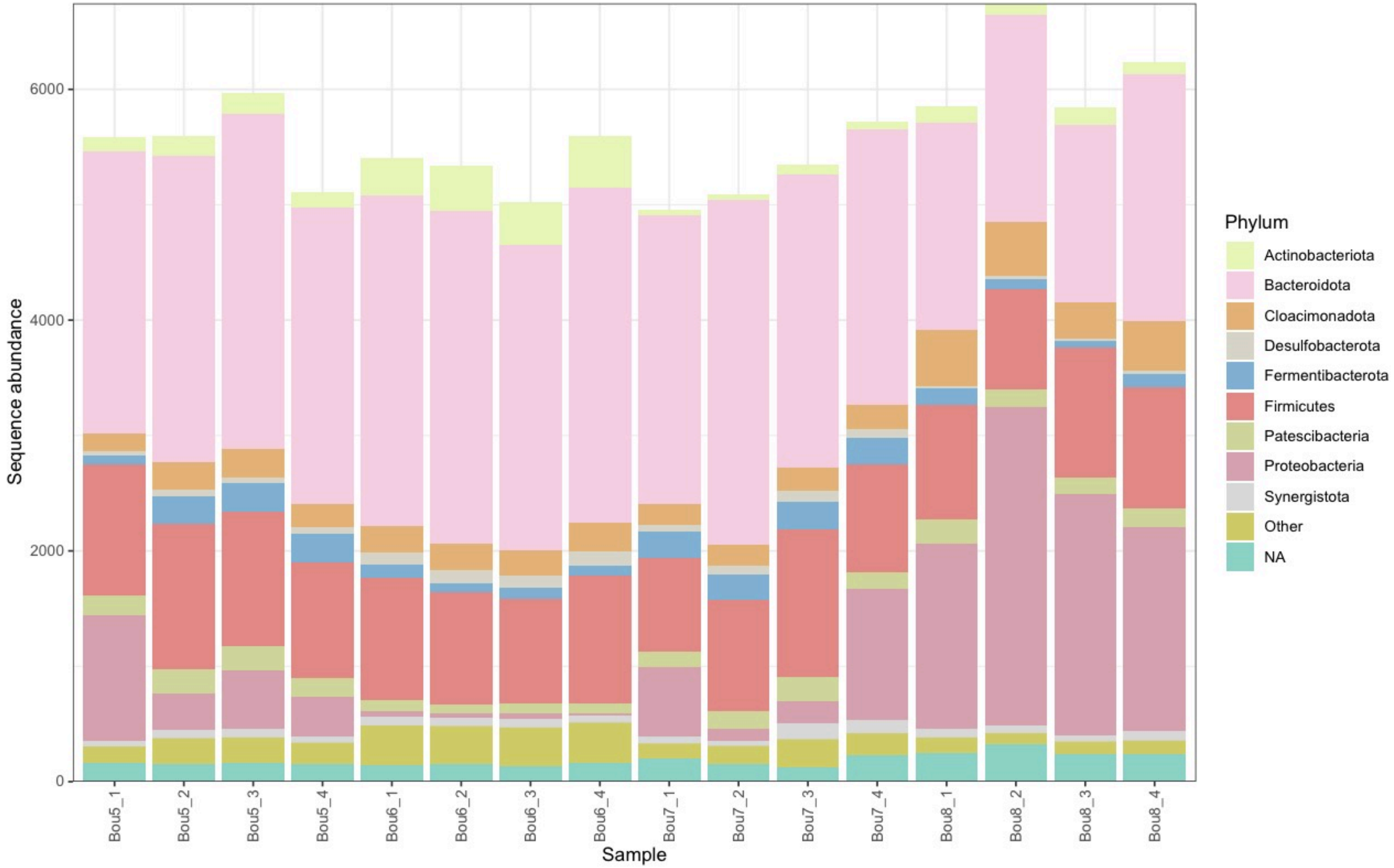

**Figure S2** B) 2022 Corn C1

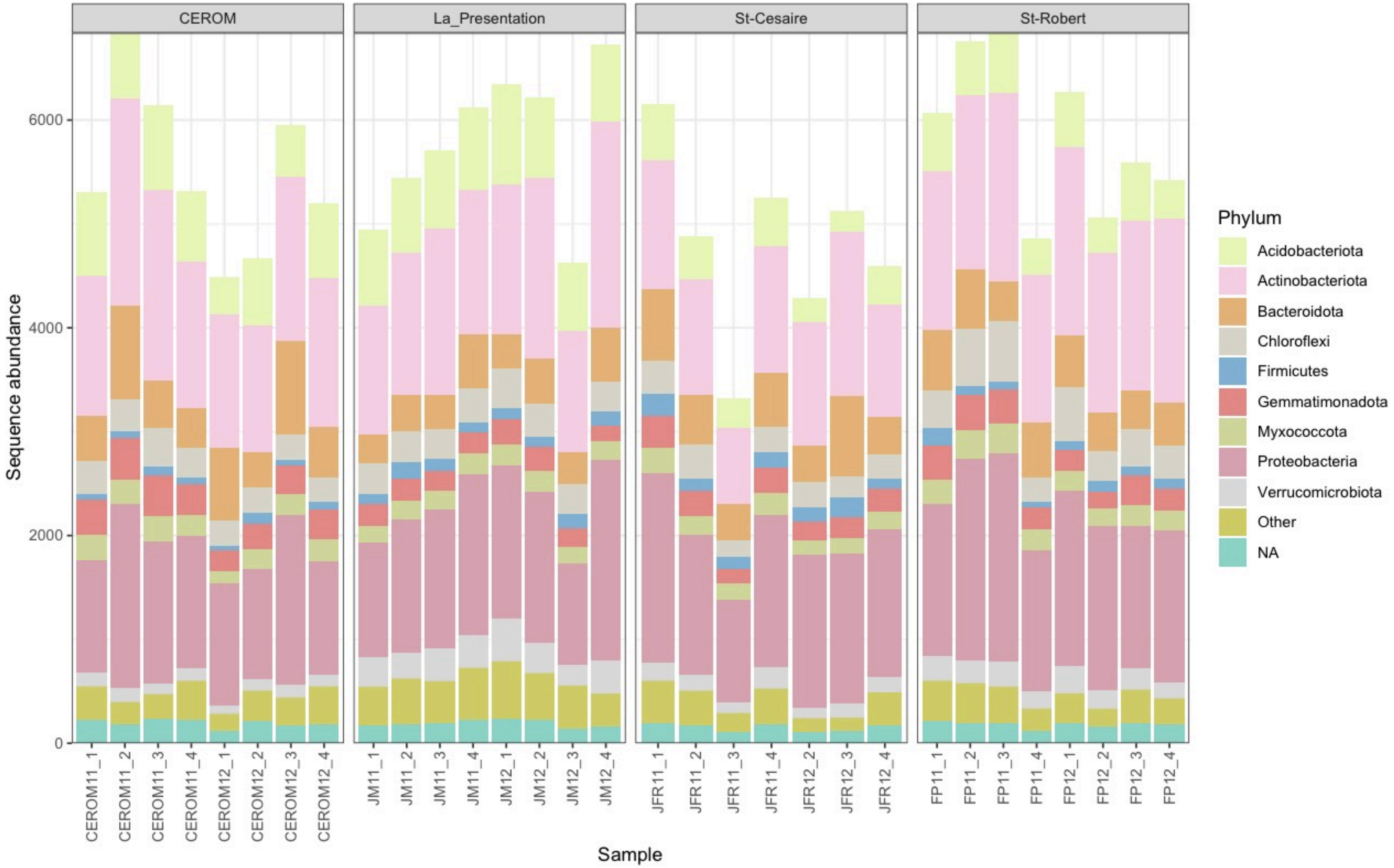

147 **Figure S2** C) 2022 Soybean C1
148

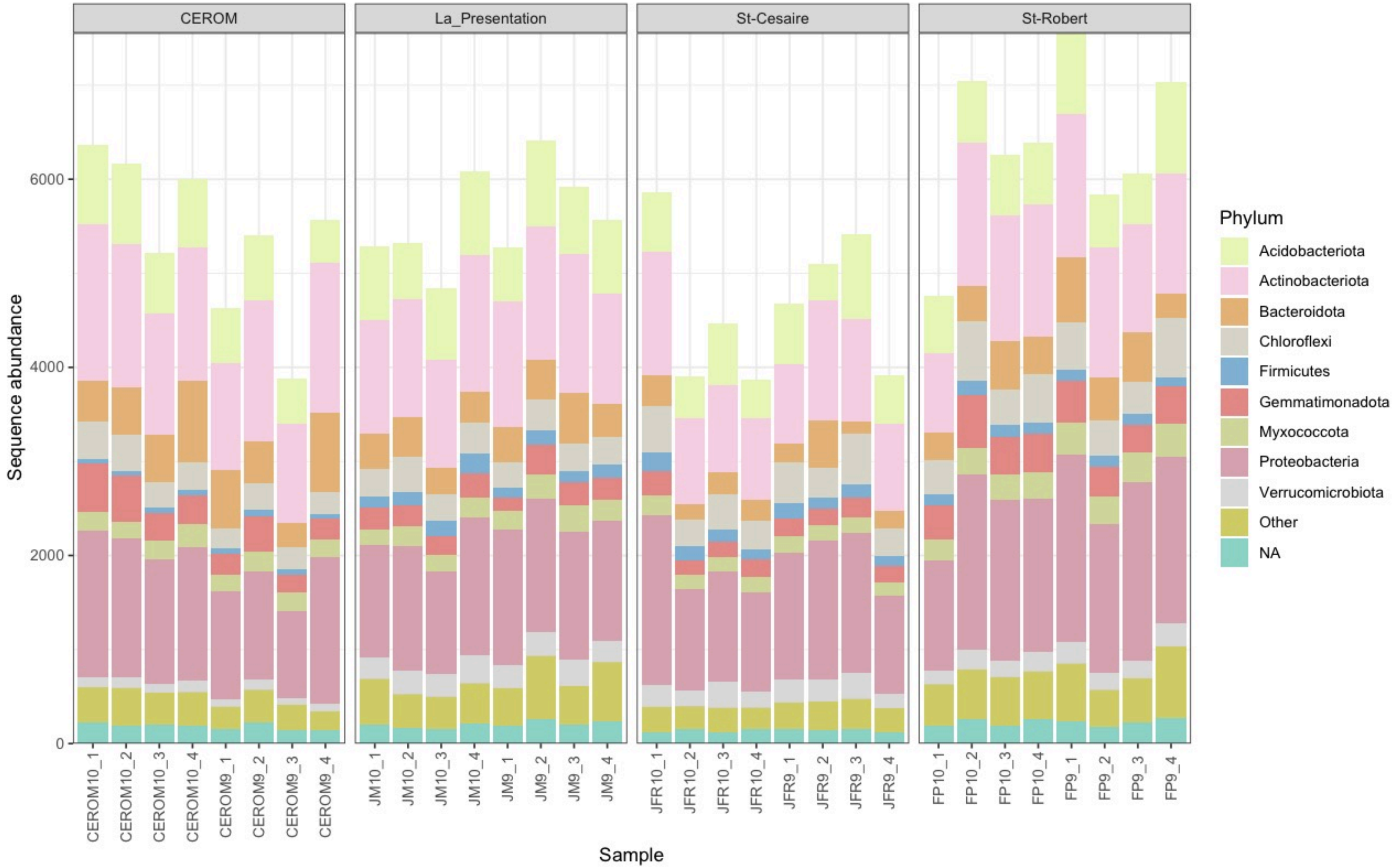


149
150

**Figure S2**. Sequence abundance of initial bacterial phyla present in each community from A) the municipal biosolids, B) corn, and C) soybean. Samples were harvested during the first campaign (C1) in May 2022 from four sites in the Montérégie area of Québec. Samples are grouped by site and colour-coded by bacterial phyla.

**Figure S3**

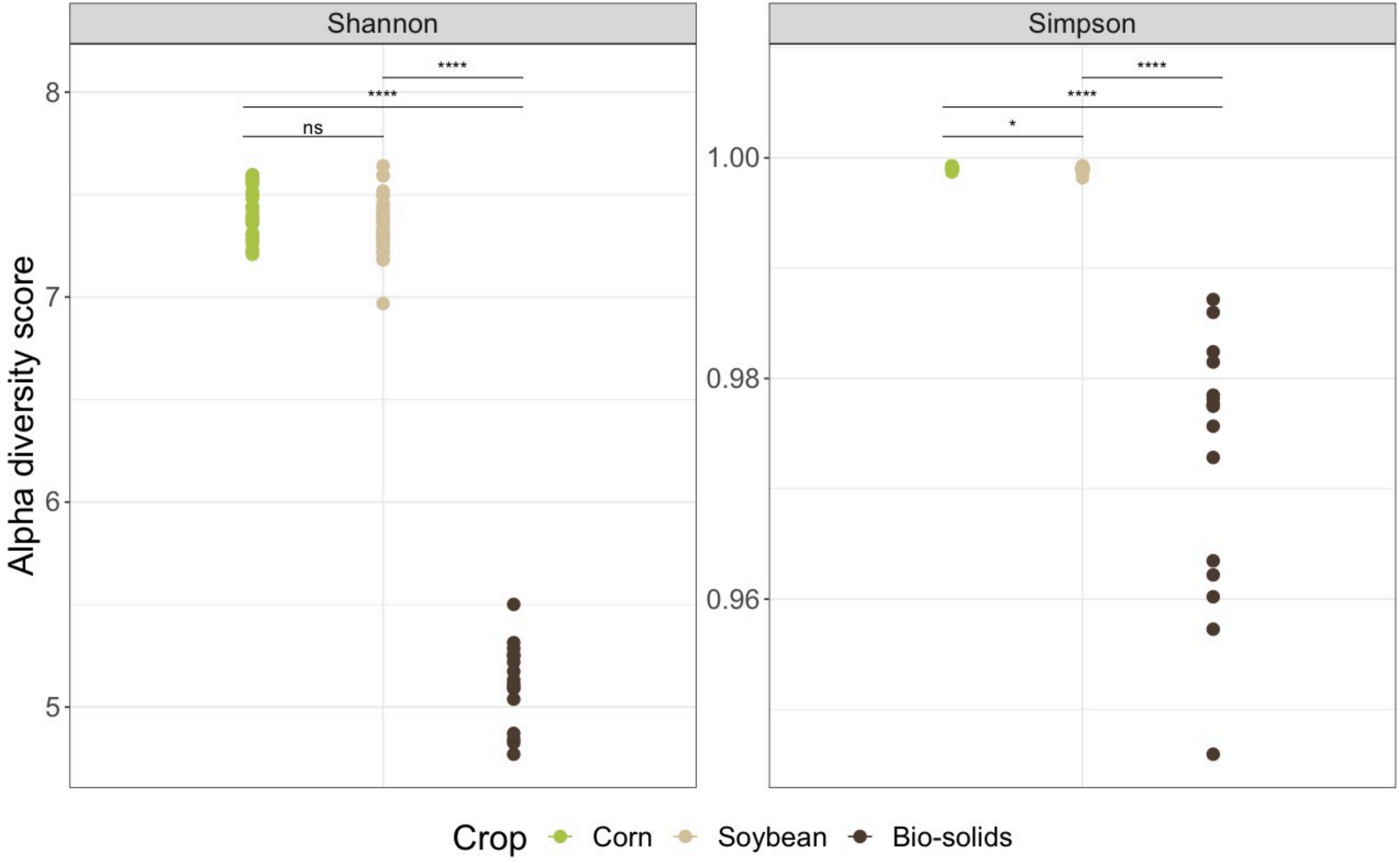
Shannon
Simpson
Alpha diversity score
8
7
6
5
1.00
0.98
0.96
ns
****
****
*
****
****
Crop
Corn
Soybean
Bio-solids

**Figure S3.** Shannon-Weaver and Simpson reciprocal diversity indices for soil bacterial communities in **2022**. Samples were harvested during the first campaign (C1, May) from four sites in the Montérégie area of Québec and bacterial composition was determined using 16S rRNA metabarcoding. MBS alpha diversity was significantly lower compared to the diversity indices of corn and soybean bacterial communities for both years (Pairwise Wilcox Test $p < 0.001$). Corn and soybean Simpson reciprocal indices are also significantly different (Pairwise Wilcox Test $p = 0.017$).

**Figure S4**

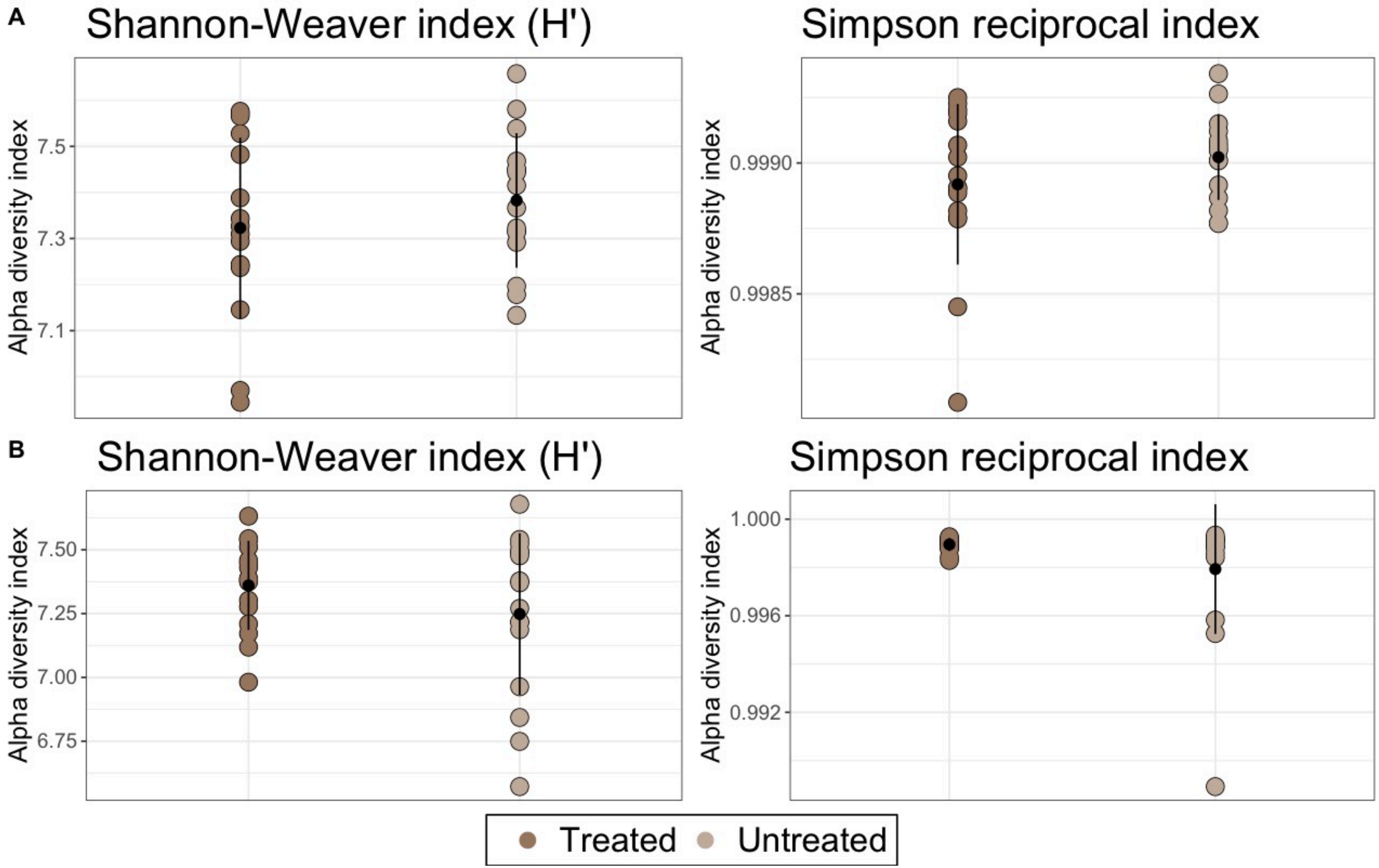
A
Shannon-Weaver index (H')
Alpha diversity index
7.5
7.3
7.1
Simpson reciprocal index
0.9990
0.9985
B
Shannon-Weaver index (H')
7.50
7.25
7.00
6.75
Simpson reciprocal index
1.000
0.996
0.992
Treated
Untreated

**Figure S4**. Shannon-Weaver and Simpson reciprocal diversity indices for soil bacterial communities from (A) corn and (B) soybean for the **2022** field trial were not significantly different ($p > 0.5$) between soil samples with and without MBS. Samples were harvested in July (C2) from four sites in the Montérégie area of Québec and bacterial composition was determined using 16S rRNA metabarcoding. A Kruskal-Wallis was used as a statistical test.

**Figure S5** A) 2021 Corn

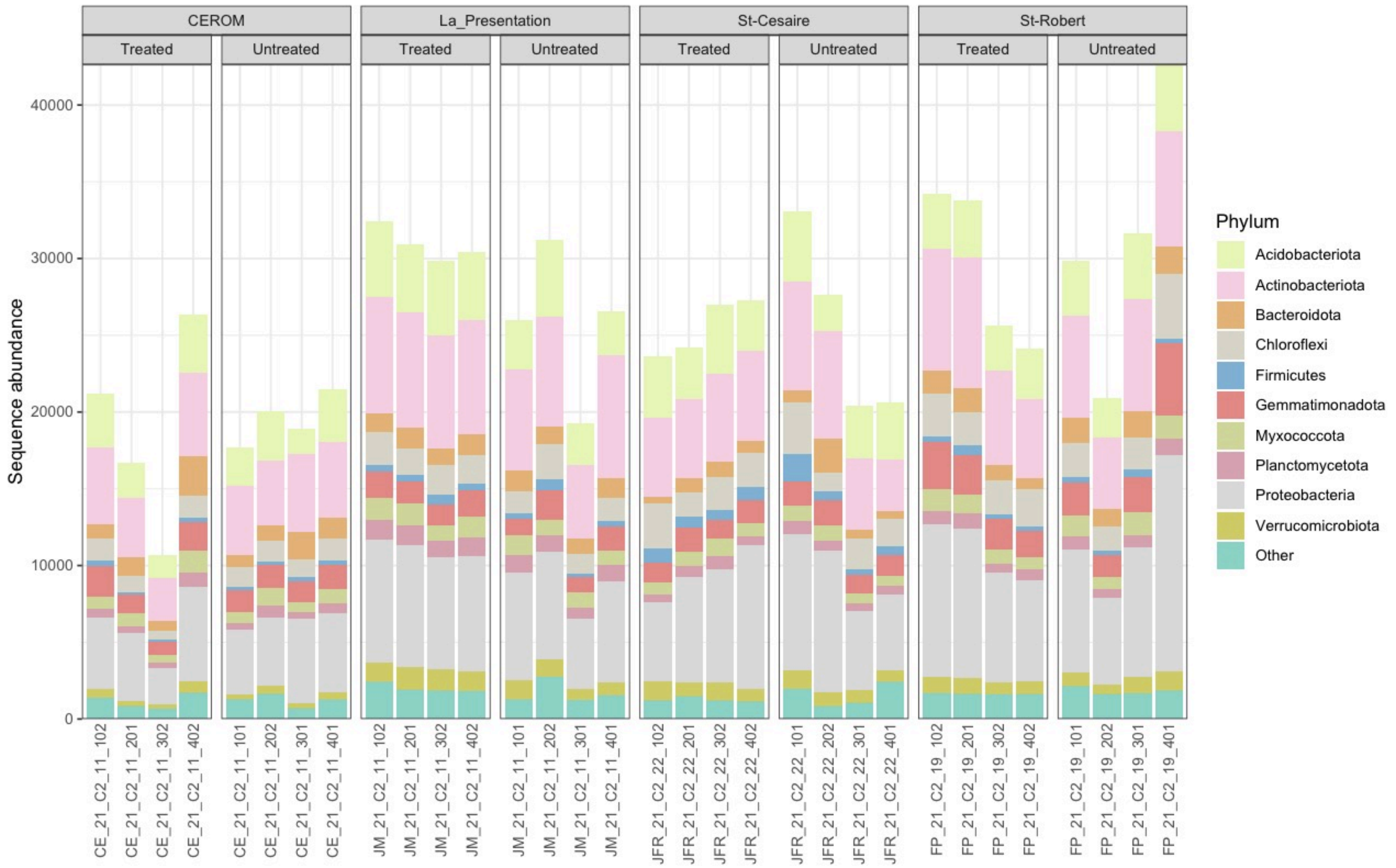

265 **Figure S5** B) 2022 Corn

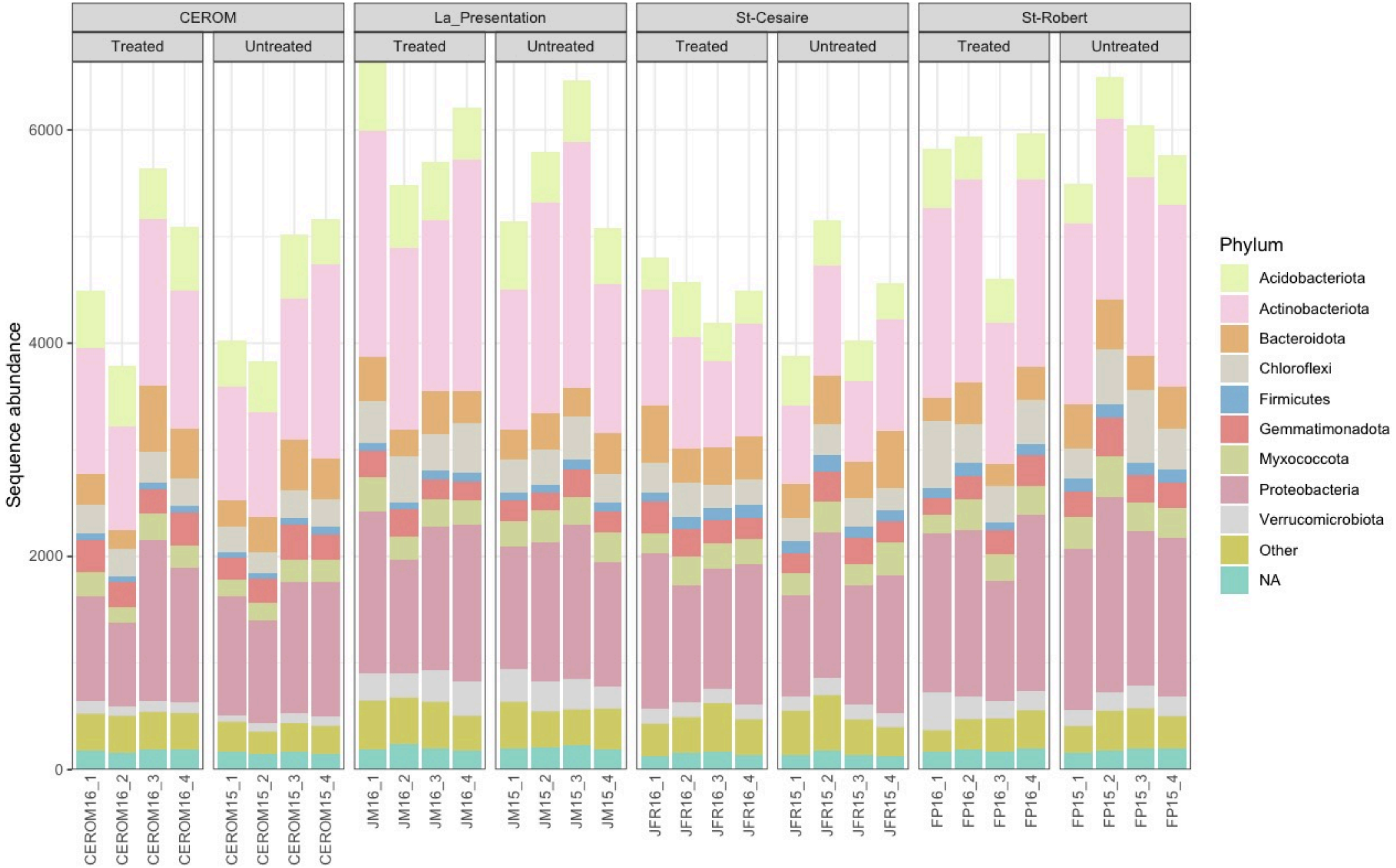

**Figure S5** C) 2021 Soybean

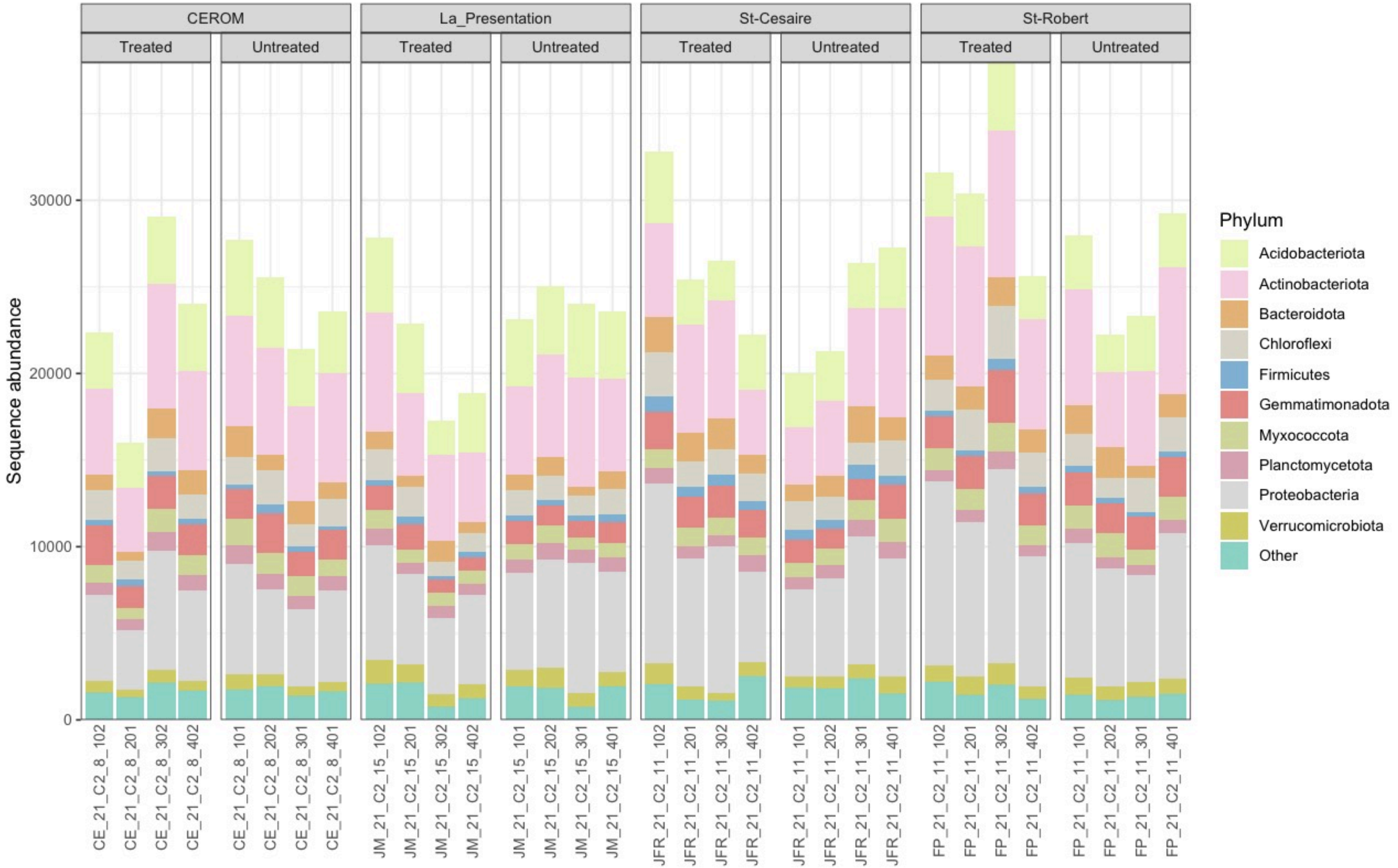

**Figure S5** D) 2022 Soybean

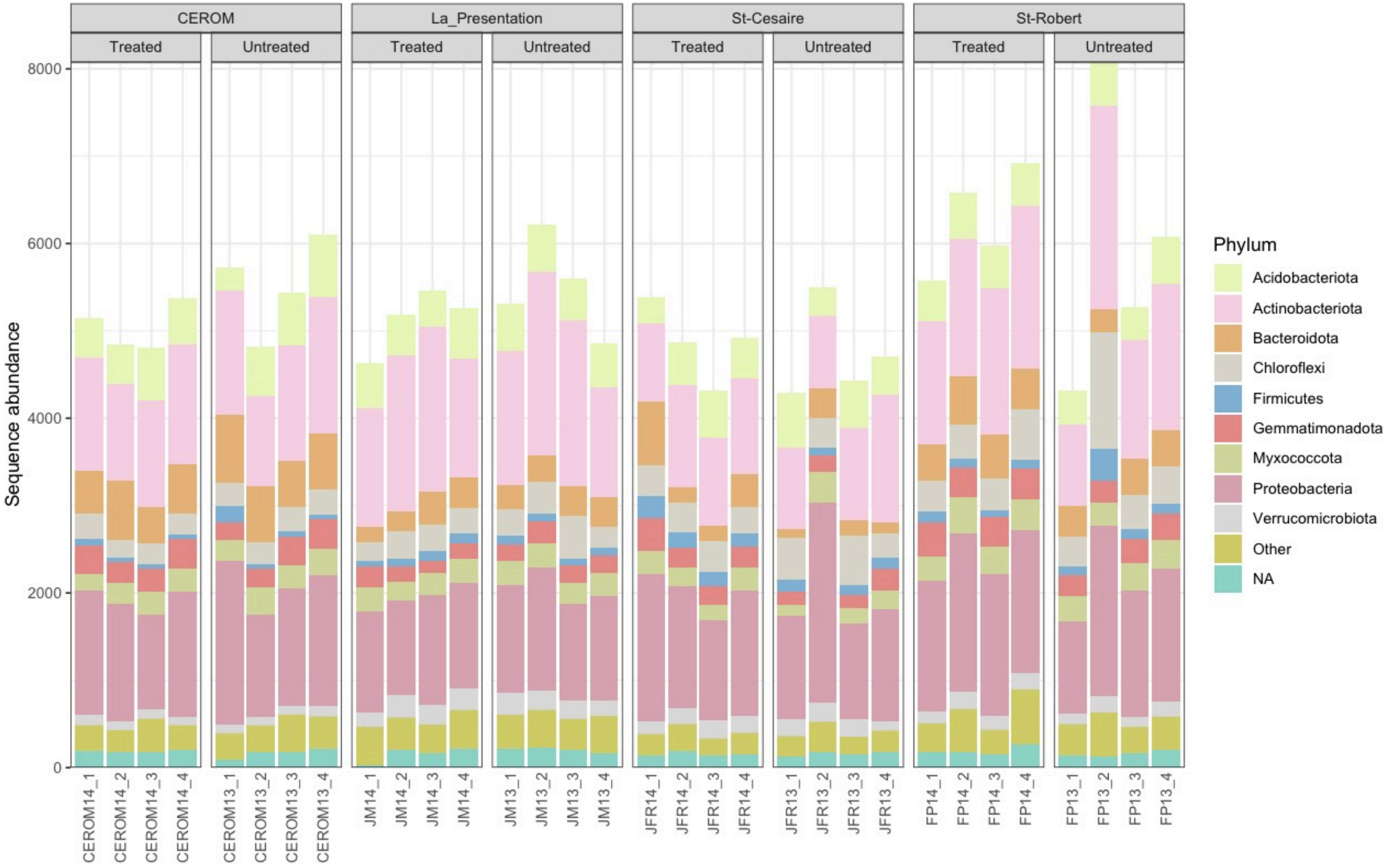

**Figure S5**. Sequence abundance of bacterial phyla present in each treated and non-treated soil community from corn (A and B) and soybean (C and D), from both trial years, 2021 (A and C) and 2022 (B and D). Samples were harvested during the second campaign (C2) in July from four sites in the Montérégie area of Québec. Samples are grouped by site and colour-coded by bacterial phyla.

**Figure S6**

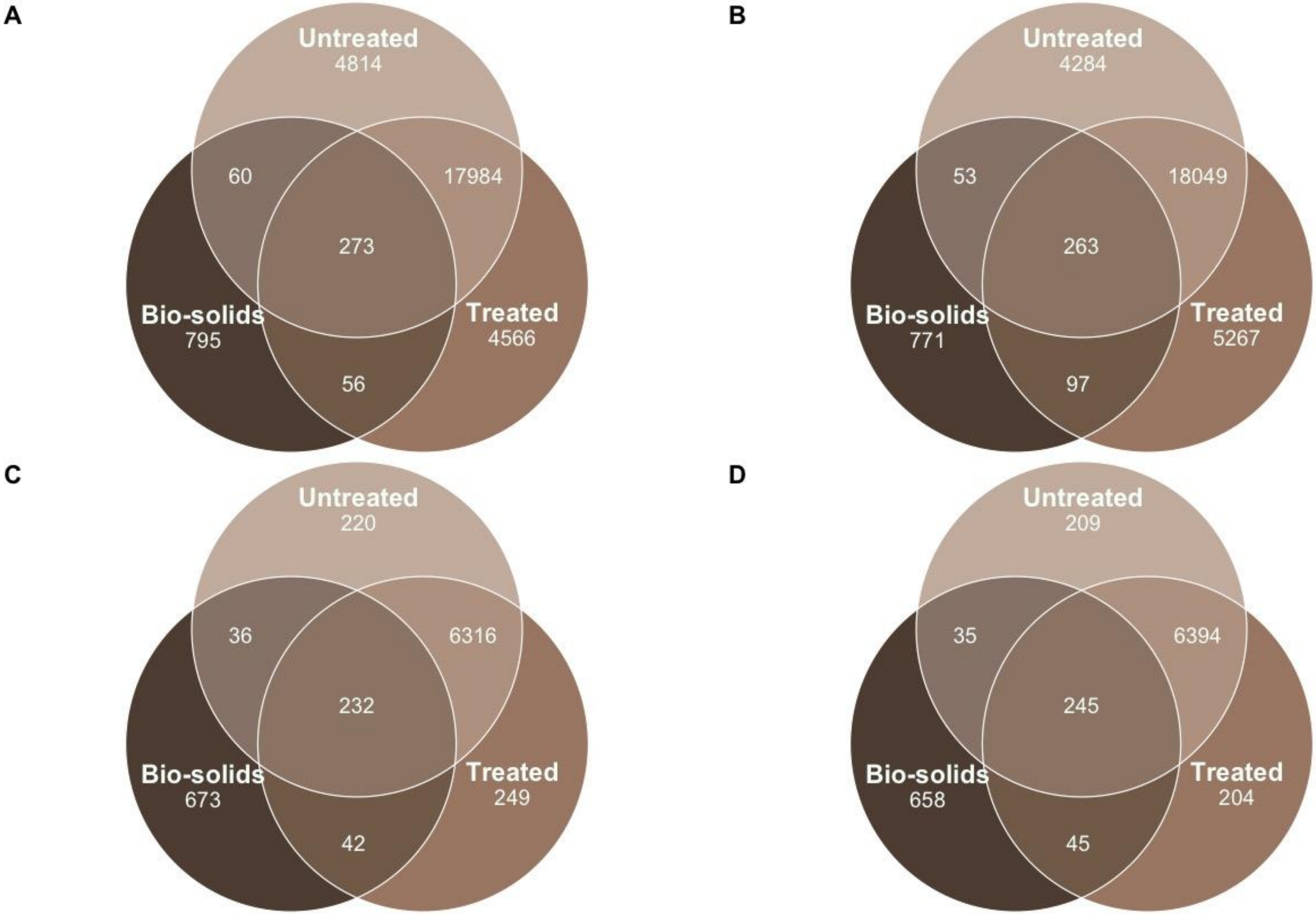
A
Untreated
4814
60
17984
273
Bio-solids
795
Treated
4566
56
B
Untreated
4284
53
18049
263
Bio-solids
771
Treated
5267
97
C
Untreated
220
36
6316
232
Bio-solids
673
Treated
249
42
D
Untreated
209
35
6394
245
Bio-solids
658
Treated
204
45

**Figure S6**. The municipal biosolids, and the MBS-treated and non-treated agricultural soils had a minority of identified ASVs in common among the bacterial communities throughout the experiment, in A) Corn 2021, B) Soybean 2021, C) Corn 2022, and D) Soybean 2022.

**Figure S7**

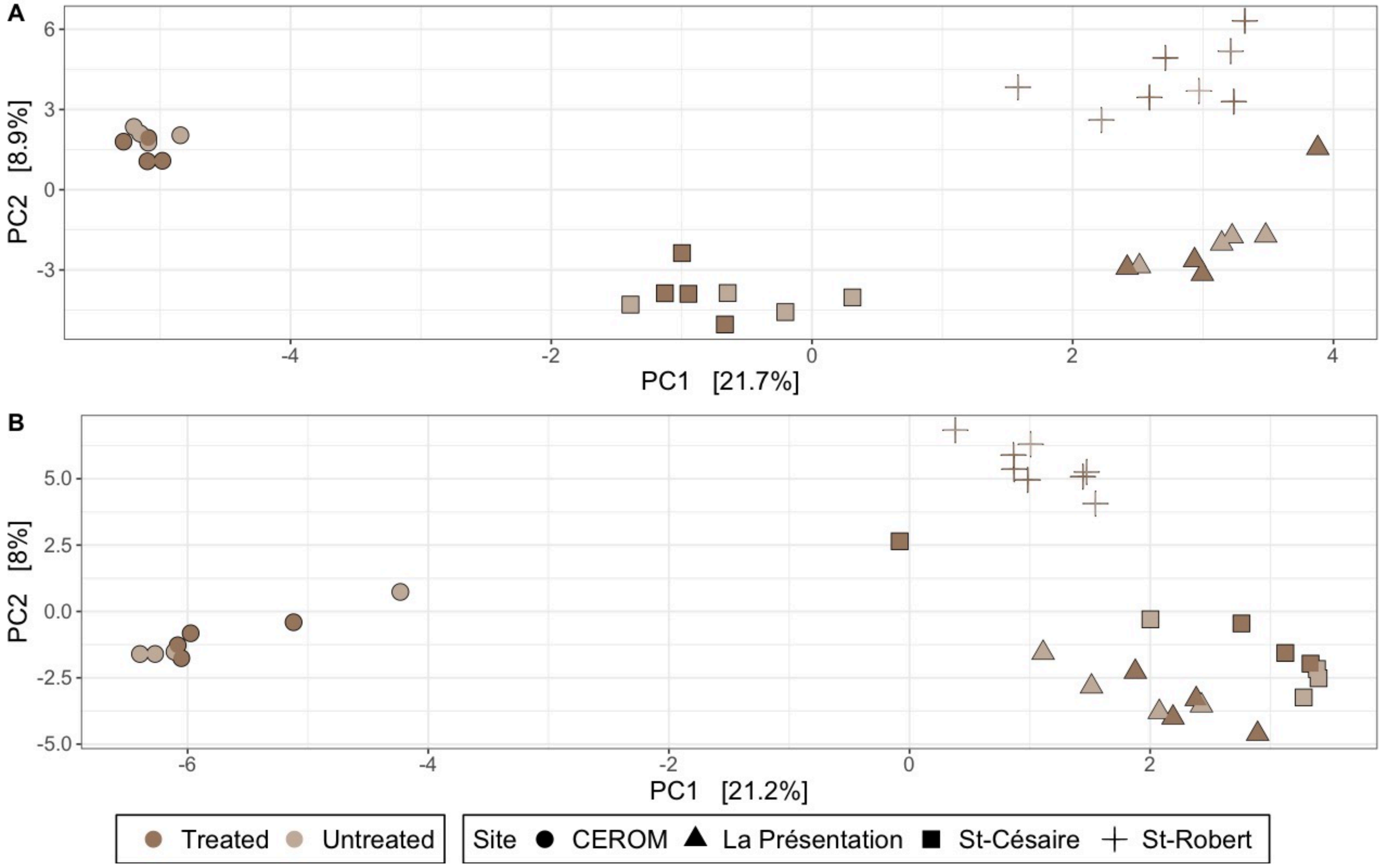
A
PC2 [8.9%]
PC1 [21.7%]
B
PC2 [8%]
PC1 [21.2%]
Treated
Untreated
Site
CEROM
La Présentation
St-Césaire
St-Robert

**Figure S7**. Soil bacterial communities treated with MBS from **2022** in corn (A) and soybean (B) were not significantly different from non-treated soils (PERMANOVA Corn: $R^2 = 0.02080$ , F = 0.9358, $p = 0.483$; Soybean: $R^2 = 0.02355$, F = 1.0139, $p = 0.359$). Samples were harvested in July (C2) from four sites in the Montérégie area of Québec. Aitchison distances appropriate for the compositional nature of sequencing data were visualised via principal component analysis. Samples are colour-coded according to their treatments and shaped according to their site.

**Figure S8** A) 2021 Soybean C1

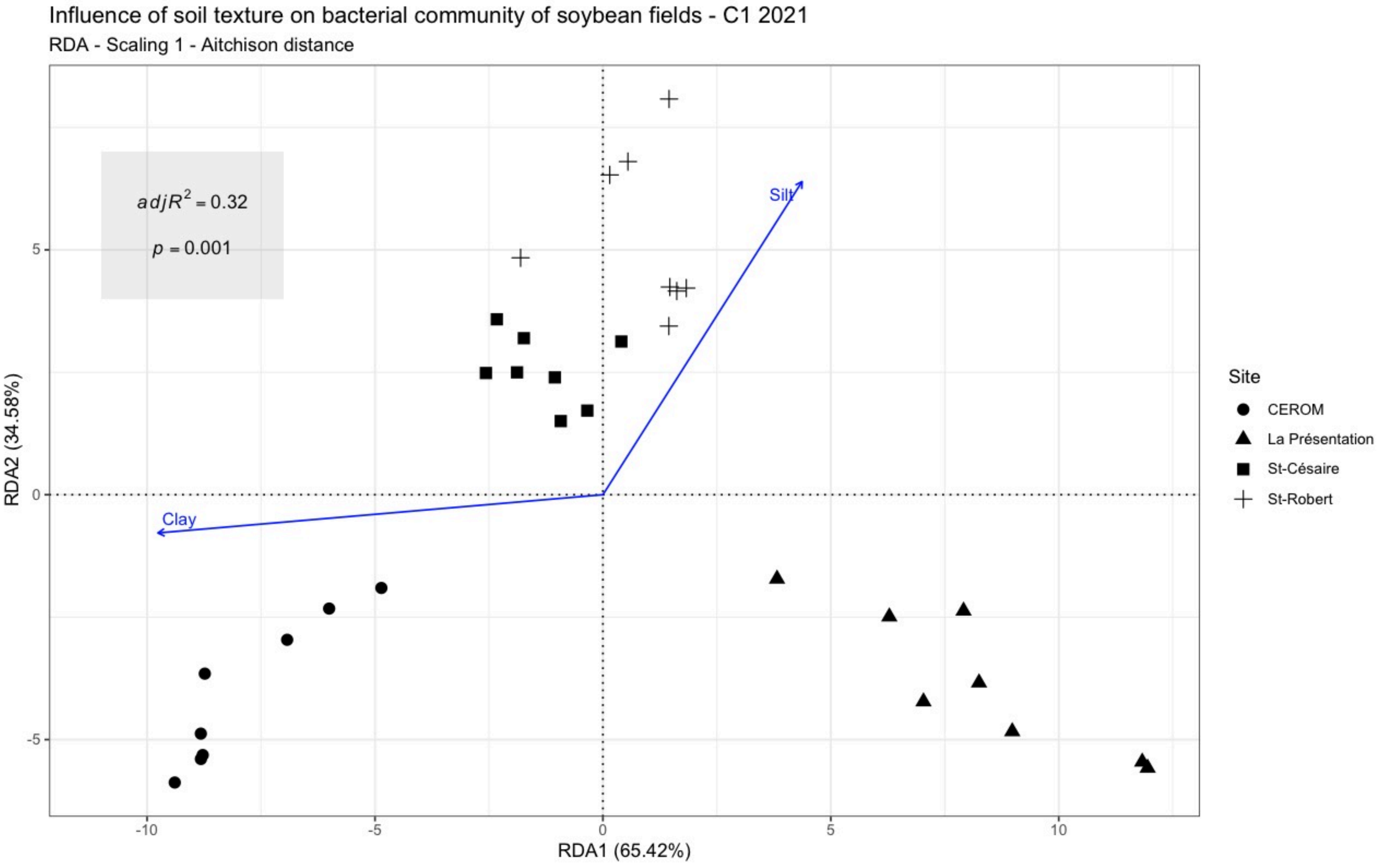

**Figure S8** B) 2021 Soybean C2

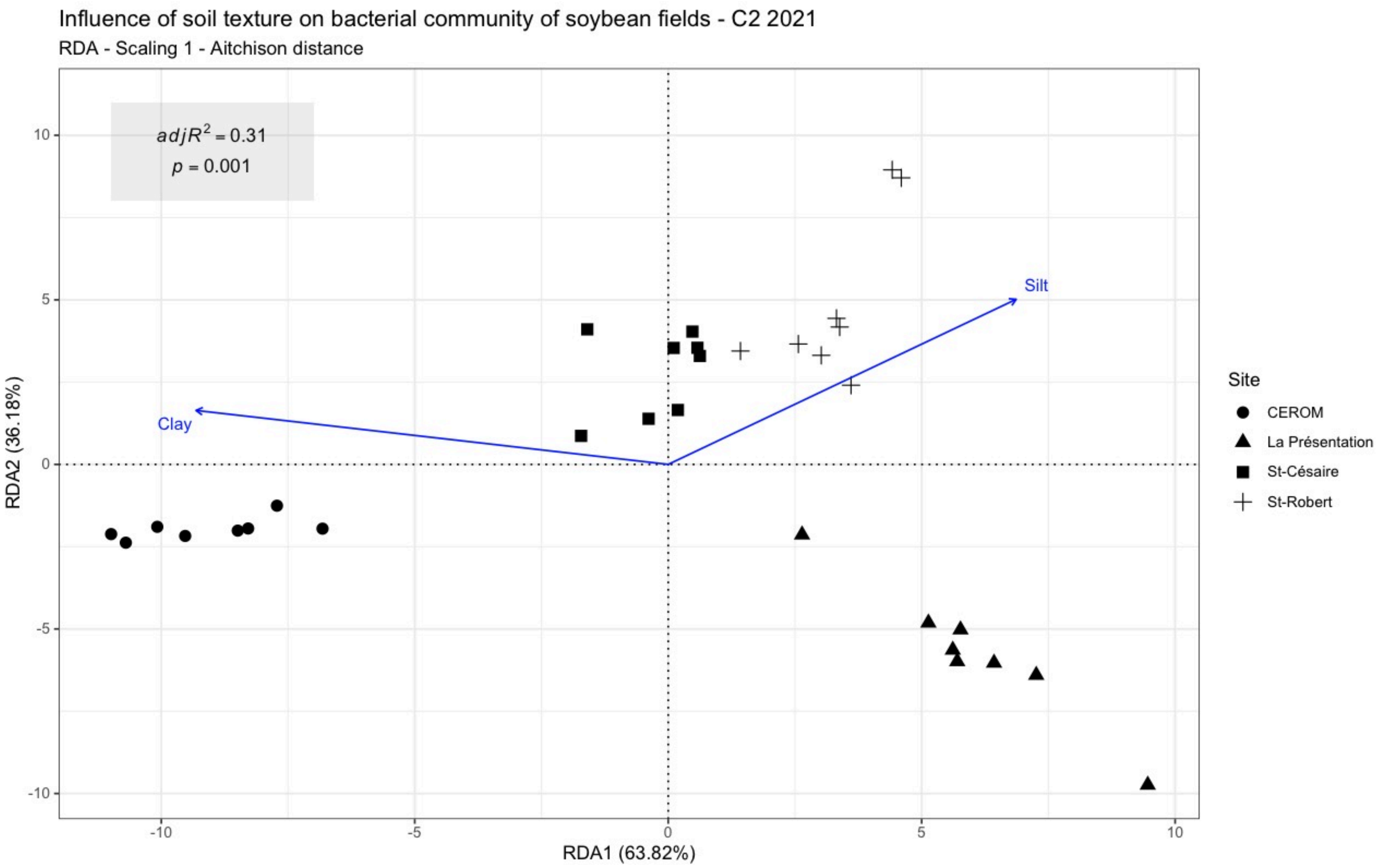

**Figure S8** C) 2022 Corn C1

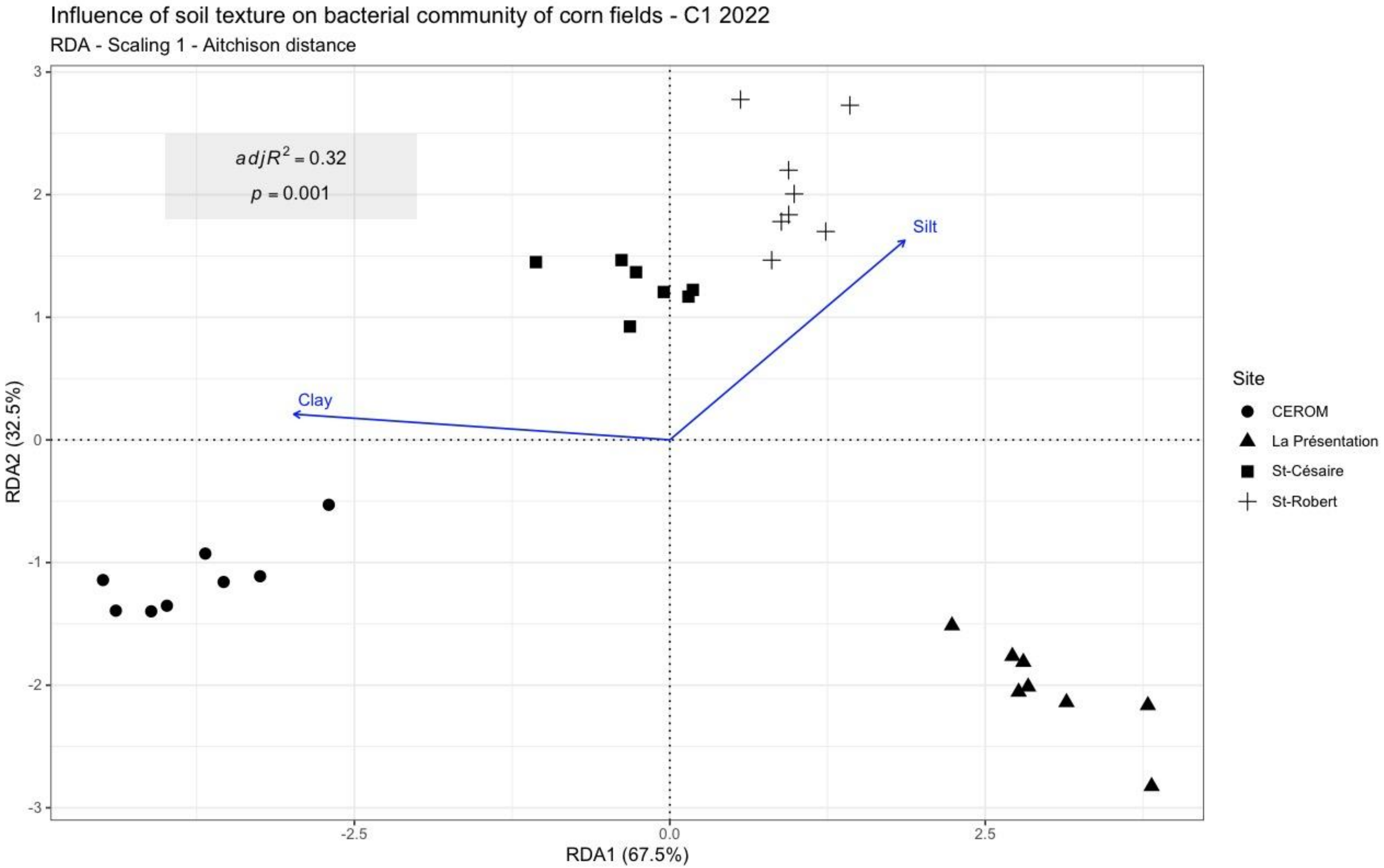

**Figure S8** D) 2022 Corn C2

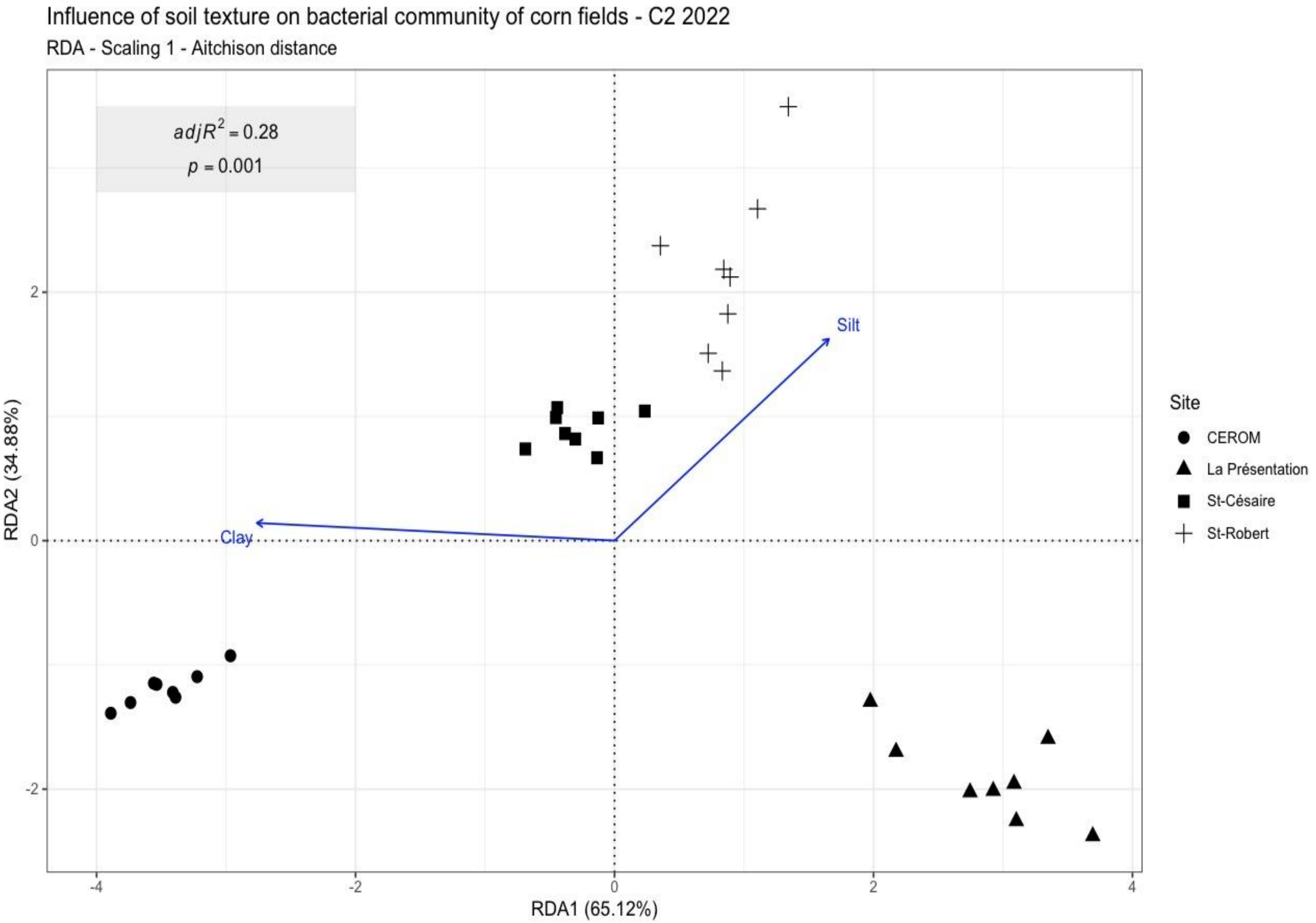

**Figure S8** E) 2022 Soybean C1

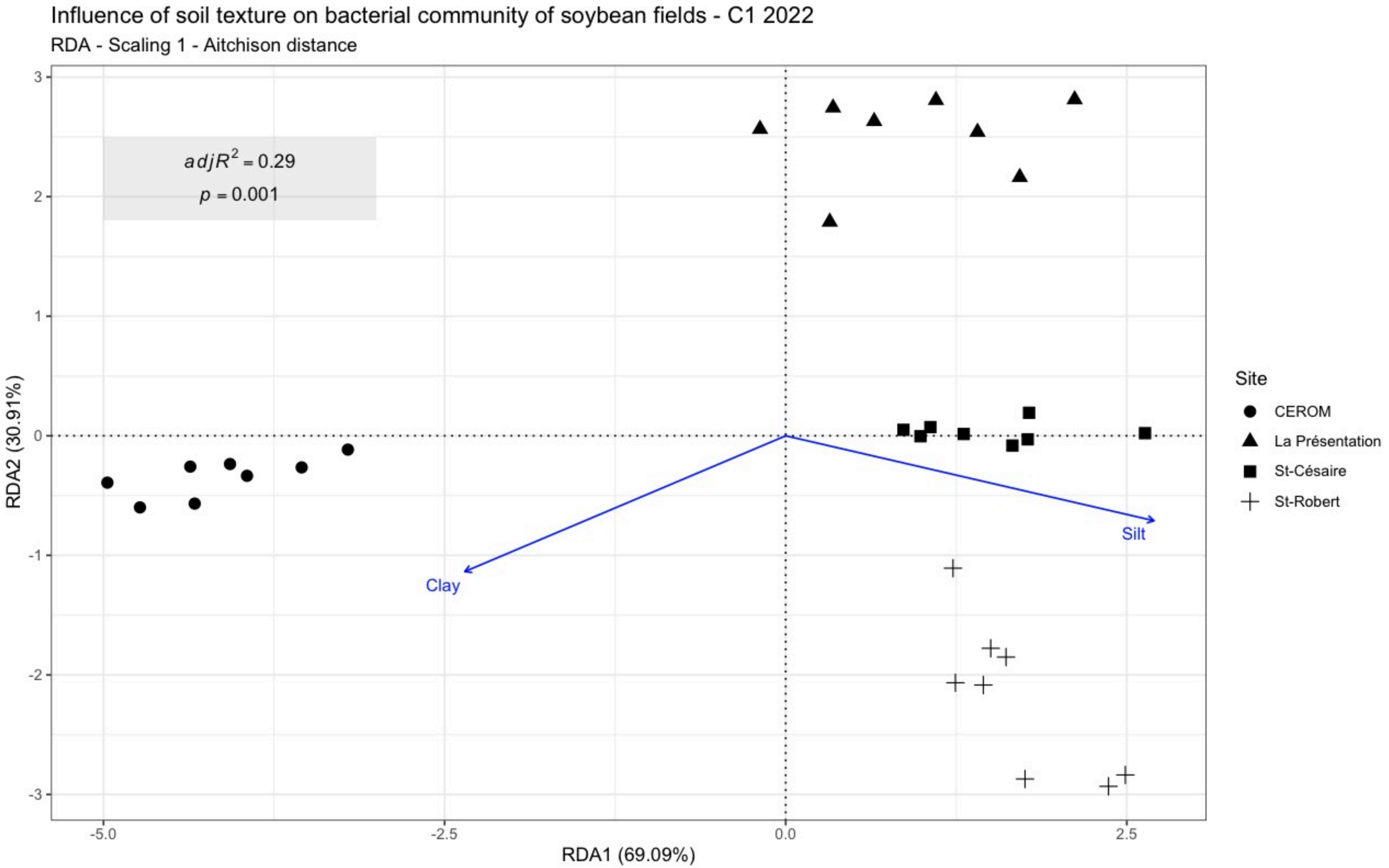

397 **Figure S8** F) 2022 Soybean C2

398

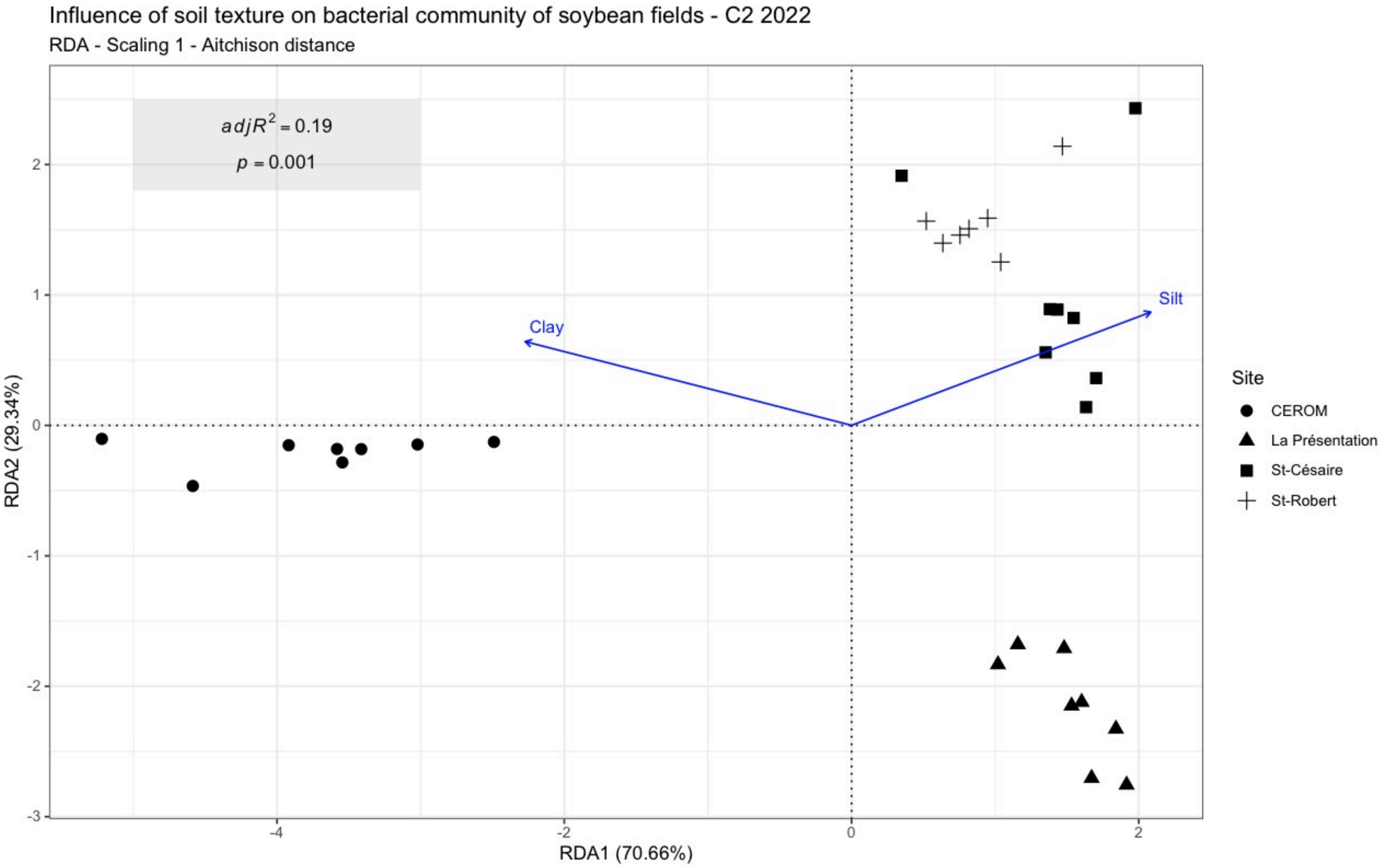


399

400

**Figure S8**. Soil bacterial communities from field experiments in corn and soybean plots were significantly structured by soil texture in 2021 (A) C1 soybean, (B) C2 soybean, and in 2022 (C) C1 corn, (D) C2 corn, (E) C1 soybean, (F) C2 soybean. One outlier sample from la Presentation was removed from the 2022 C2 soybean analysis. Samples were harvested in May (C1) and July (C2) from four sites in the Montérégie area of Québec. An Aitchison distance matrix, appropriate for the compositional nature of sequencing data, was used in the distance-based redundancy analysis. The RDA quantified how soil texture structured the bacterial communities, where those with similar ASV composition were plotted closer together. Soil bacterial communities from each sample are represented by different shapes for each site.

**Figure S9**

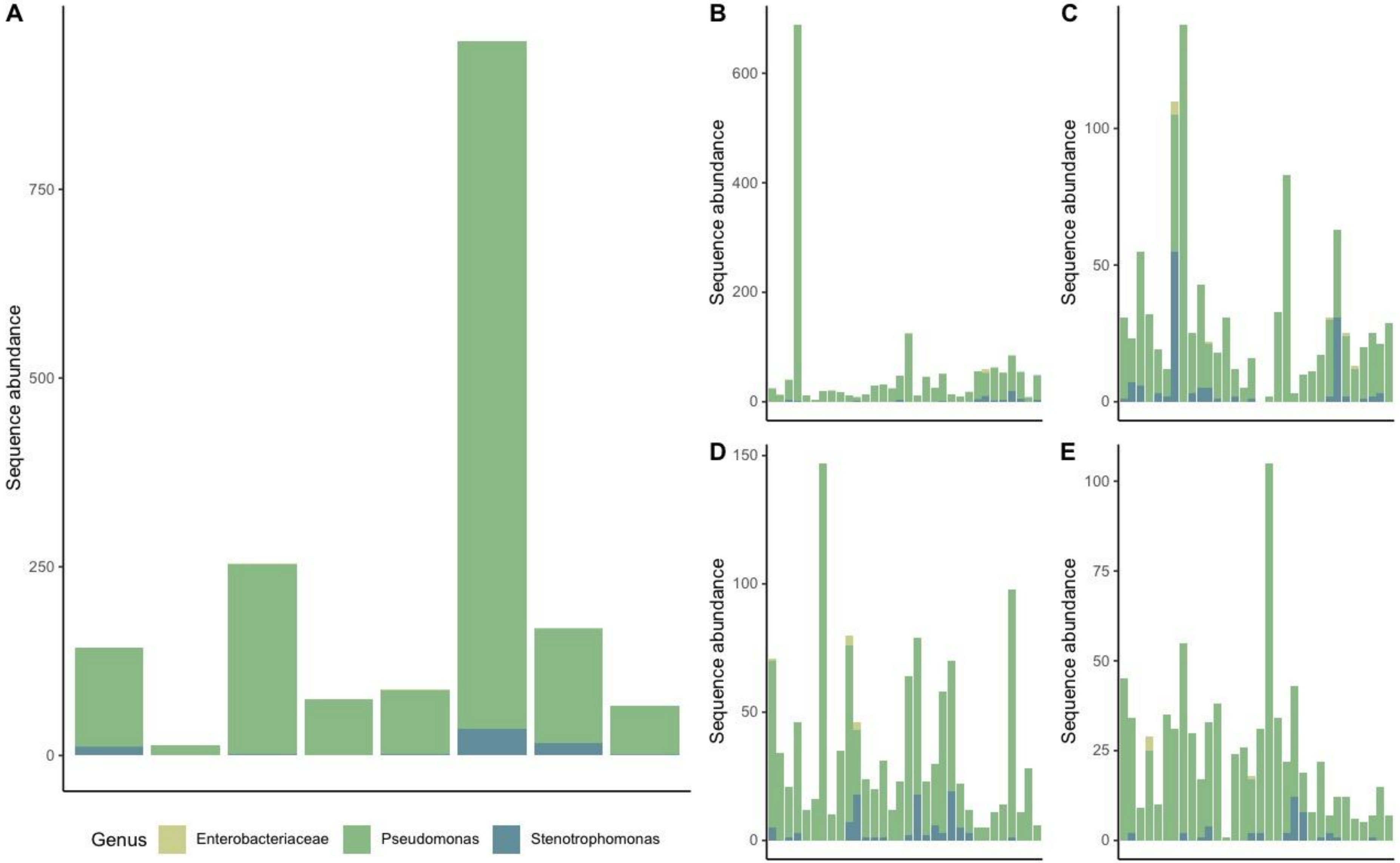
A
B
C
D
E
Sequence abundance
0
250
500
750
200
400
600
50
100
150
25
75
Genus
Enterobacteriaceae
Pseudomonas
Stenotrophomonas

Figure S9 con't

**Figure S9** con't

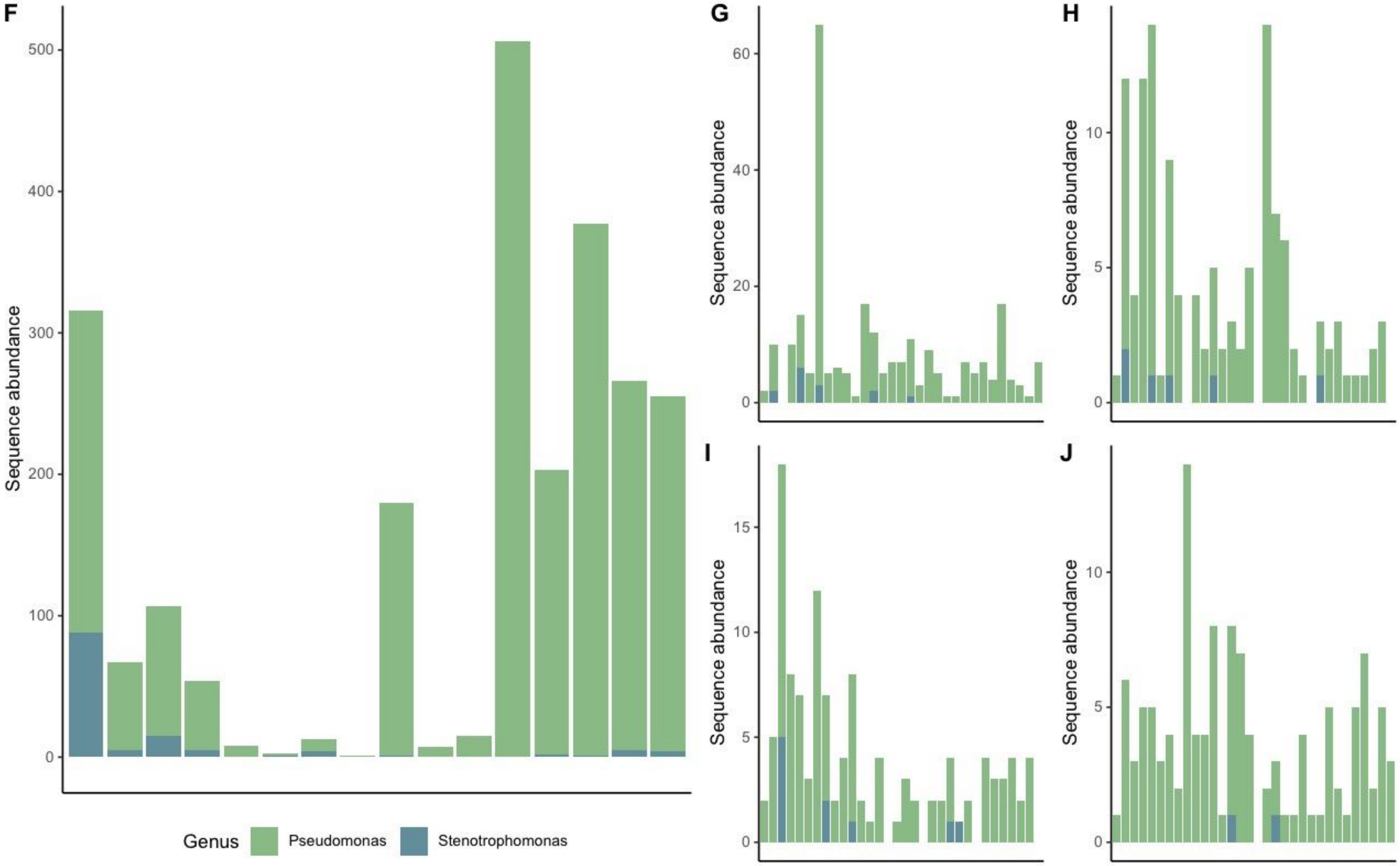

**Figure S9**. Abundance of 16S rRNA sequences associated with potential human bacterial pathogens in municipal biosolids and in the agricultural soils: A) MBS 2021, B) corn 2021 C1, C) corn 2021 C2, D) soybean 2021 C1, E) soybean 2021 C2, F) MBS 2022, G) corn 2022 C1, H) corn 2022 C2, I) soybean 2022 C1, J) soybean 2022 C2. Bar plots represent the samples harvested during 2021 (A-E), and 2022 (F-J), at two timepoints (C1, May; C2, July), over the growing seasons from four sites in the Montérégie area of Québec. Different bacterial genera detected in each sample are colour-coded
